# Huntington’s Disease Produces Multiplexed Transcriptional Vulnerabilities of Striatal D1-D2 and Striosome-Matrix Neurons

**DOI:** 10.1101/2022.04.25.489455

**Authors:** Ayano Matsushima, Sergio Sebastian Pineda, Jill R. Crittenden, Hyeseung Lee, Kyriakitsa Galani, Julio Mantero, Manolis Kellis, Myriam Heiman, Ann M. Graybiel

**Author notes:** Correspondence (AMG), (MH), (MK). These authors contributed equally to this work (Ayano Matsushima, Sergio Sebastian Pineda).

## Abstract

Striatal cell-type-specific vulnerability in Huntington’s disease (HD) preferentially affects dopamine D2R-expressing projection neurons (SPNs), compatible with manifest motor symptomatology in HD. Transcriptional studies of striatal striosome-matrix compartmentalization in HD are, however, limited, despite pathologic evidence for striosome vulnerability aligning with early mood symptomatology. We used single-nucleus RNA-sequencing on striatal samples from two murine models, and rare Grade 1 HD patient tissues, to examine striosome and matrix sub-clusters within parent D1 and D2 SPN clusters. In human HD, striosomal SPNs were the most depleted SPN population. Surprisingly, for both mouse models, transcriptomic distinctiveness was diminished more for striosome-matrix SPNs than for D1-D2 SPNs. Compartmental markers were dysregulated so as to cancel endogenous identities as striosomal or matrix SPNs, but markers for D1-D2 exhibited less identity obscuring. The canonical striosome-matrix as well as D1-D2 organizations of the striatum thus are both strongly, but differentially, compromised in HD and are targets for therapeutics.

## Introduction

Huntington’s disease (HD) is a major extrapyramidal disorder typically characterized by early-stage mood disorders, a subsequent hyperkinetic and/or hypokinetic ‘manifest’ stage, and an eventual decline to death^1^. Expansion of uninterrupted CAG repeats within the mutant *HTT* gene (*mHTT*) reaching over 40 results in manifest HD. A hallmark of HD is the profound loss of neurons in the neostriatum. Work on HD models and human HD brain samples has documented marked anatomic and electrophysiologic alterations within the striatum. Striatal spiny projection neurons (SPNs) expressing D2 dopamine receptors and giving rise to the indirect pathway of the basal ganglia (iSPNs) are differentially vulnerable^2–5^. With time, multiple striatal cell types become affected, including the direct pathway SPNs expressing D1 receptors (dSPNs) and even glial cells, leading to cavitation of the striatum. These pathophysiologic patterns are concordant with the hyperkinetic motor symptomatology in HD, as the strongly perturbed iSPNs normally support motor inhibition.

A second, less fully studied feature of striatal vulnerability has been found to involve the neurochemical compartmental organization of the striatum, in which molecularly specialized labyrinthine ‘striosomes’ wind through the surrounding matrix compartment^6^. Both striosomes and matrix contain both dSPNs and iSPNs^7–10^, and, like the D1-D2 axis of striatal organization, the striosome-matrix (S-M) axis specifies input-output connectivity patterns. The fact that anteromedial striosomes favor limbic circuits and much of the matrix favors sensory-motor circuits^11, 12^ has raised interest in the possibility that dysregulation along the S-M axis could be related to the modal transition of HD symptomatology over time. Reports of early vulnerability of striosomes based on post-mortem anatomy^13, 14^, especially in identified mood-disorder patients^15^, have led to the view that striosomal dysfunction could differentially contribute to the pre-manifest periods, with mood disorders, then merge with following motor dysfunction as the matrix becomes increasingly affected^14, 16^.

Transcriptomic studies have since also indicated differential vulnerability of the striosomes and matrix in HD. Substance P/Tac1, a marker of striosomes, was found to be downregulated especially in dSPNs, suggestive of an intensified loss of markers of the striosome compartment^17^. However, it is not yet clear how transcriptional dysregulation in SPNs along the S-M organizational axis of the striatum relates to the dysregulation of the seemingly orthogonal D1-D2 (dSPN-iSPN) axis of organization. Here, to address this issue, we leveraged single-nucleus RNA-sequencing (snRNA-seq) to examine striatal transcripts derived from the human HD striatum, and further compared in detail striatal transcripts from two mouse models of HD, the classic R6/2 model with rapid progression, and the knock-in zQ175 model with slower progression. For the human as well as rodent transcripts, we applied advanced sub-clustering of dSPN and iSPN populations and used a curated set of S-M marker genes to annotate striosome- and matrix-selective sub-clusters inside them^18, 19^. With this base, we then compared the differential transcriptomic changes in the cell types according to their compartmental sub-clusters and by their D1-D2 parent clusters.

Our evidence demonstrates that, despite the dominant dysregulation of D2-expressing iSPNs as widely accepted, the transcriptomic profiles differentiating striosomal SPNs from matrix SPNs were more dysregulated than those distinguishing D1 SPNs from D2 SPNs. Transcripts differentiating striosomal and matrix SPNs, classified as S-M ‘markers’ in our analysis, were dysregulated in a cell-type-specific manner so as to blur the endogenous transcriptional distinctiveness of the two compartments. Striosomal SPNs exhibited upregulated matrix markers but downregulated striosome markers, thus diminishing their striosome-like identity. Matrix SPNs exhibited upregulated striosome markers, but downregulated matrix markers, thus diminishing their matrix-like identity. In sharp contrast, D1-D2 marker transcripts were dysregulated irrespective of cell type. Thus, both mouse models of HD exhibited a cohesive pattern tending to cancel out the endogenous identities of striosome and matrix SPNs, whereas the distinction along the D1-D2 (dSPN-iSPN) axis was more robust despite the greater D2 vulnerability, as confirmed in the human Grade 1 case.

In both HD mouse models, we also found that the absolute degree of cell-type-specific dysregulation of gene expression, whether up or down, was most prominent in a particular newly identified set of putative iSPNs that we classified as ‘outlier-D2’ (O-D2), followed by striosomal iSPNs (S-D2). The O-D2 SPNs formed a small sub-cluster within the parent D2 cluster. This pattern accords with the well-known D2-dominant deficits in HD and was consistently observed in dysregulations of all genes including non-markers, meeting the criterion of being cell-type-specific (i.e., dysregulated in opposite directions in different cell types). It was for the genes that were dysregulated in the same directions across cell types that the two HD models exhibited unique transcriptomic alterations, likely reflecting their distinct genetic makeups. The severe loss of SPNs in HD patients, which was not found in HD murine models, hindered in-depth transcriptomic analysis in human samples. Yet, we were able to analyze a Grade 1 HD case and found cell-type-specific vulnerability in the pattern of depletions: S-D2 were the most severely depleted SPNs of the entire SPN population, followed by S-D1 and M-D2.

These findings suggest that transcriptomic vulnerability in HD is constrained not only by the canonical D1-D2 pathway differentiation of SPNs, but also by their compartmental striosome-matrix distinctions. This result suggests a decidedly multiplexed order of SPN-type-specific vulnerability in the striatum in HD that could differentially contribute to the pre- manifest and manifest stages of this severe basal ganglia disorder.

## Results

We analyzed snRNA-seq data by using ACTIONet^20–34^ (Extended Data Fig. 1), from striatal samples harvested from human striatum and from R6/2 and zQ175 HD model mice, and originally reported in an initial study without attention to the coordinated compartmental transcriptomics examined here (Fig. 1A-C)^17^. We further collected data from a rare Grade 1 case, including from samples of both the caudate nucleus and the putamen, newly reported here. We collected 62,487 nuclei across twelve controls, and the Grade 1 HD case. For the mouse models, the numbers were 112,295 nuclei across fifteen mice: eight isogenic control and seven R6/2 model mice; and 63,015 nuclei across eight mice: four isogenic control and four zQ175DN model mice (Supplementary Table 1). R6/2 (and their control, CBA) mice were harvested at 9 weeks of age, and zQ175 (and their control, BL6) mice at 6 months of age. They were taken to represent an early- to mid-phenotypic time point of each model, based upon whole-tissue RNA sequencing studies^4^.

**Fig. 1.**
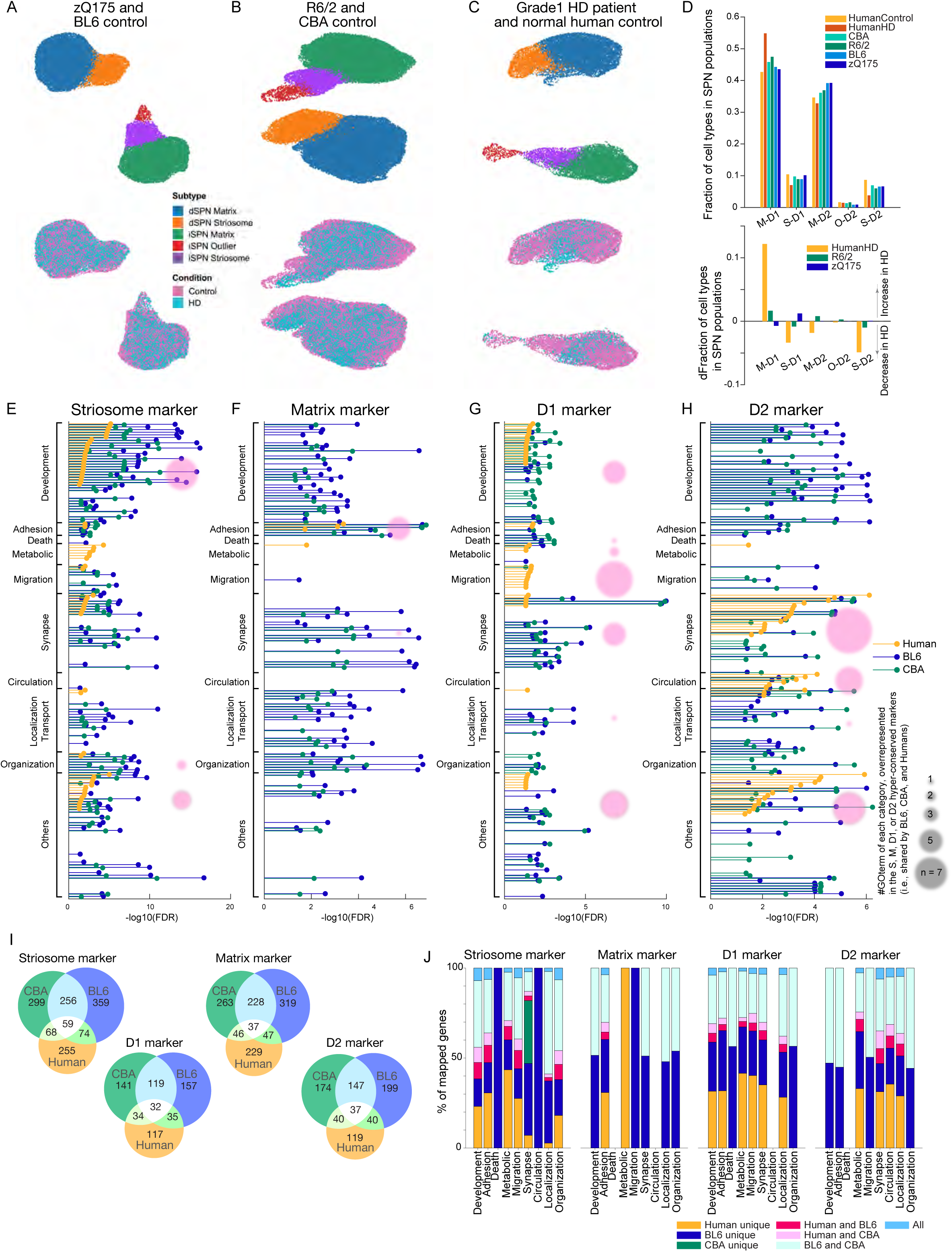
Identification and characterization of human and rodent SPN cell-type-specific markers. **A-C,** ACTIONet UMAP plots of distinct dSPN (D1) and iSPN (D2) parent clusters, and striosomal and matrix sub-clusters within zQ175 and their control samples (**A**), R6/2 and their control samples (**B**), and a human Grade 1 HD patient and healthy controls (**C**). Samples described in Supplementary Table 1. **D**, Fraction of each SPN subtypes in the entire SPN populations identified (top) and the change of fractions in HD as compared to controls (bottom). **E**, FDR for enriched GO terms in universal striosome markers found in BL6 (blue), CBA (green), and humans (orange). Within the GO terms overrepresented in either of striosome, matrix, D1, or D2 markers, top (i.e., lowest FDR) 40 GO terms are included, and grouped into 9 categories. In each group, the GO terms are sorted by FDRs for humans. Pink circles indicate the number of GO terms in each category, found to be commonly overrepresented in the hyper-conserved striosome markers. **F-H**, Same as *E* but for universal matrix (**F**), D1 (**G**), or D2 (**H**) markers. **I**, Venn diagrams showing the marker overlaps across species or mouse strains for universal striosome, matrix, D1 or D2 markers as labeled. **J**, Overlap of marker genes across species that are mapped onto GO terms related to development, adhesion, metabolic, migration, synapse, circulation, localization/transport, and organization (see Supplementary Table 2 for GO IDs included). See also Extended Data Figs. 1-5.

With a curated set of markers (Supplementary Table 1, Extended Data Fig. 2), we identified striosomal (S) and matrix (M) SPN sub-clusters within both the D1 and the D2 clusters in the mice (Fig. 1A,B), as previously described^18, 19^, as well as in humans (Fig. 1C). In addition to S and M clusters, we identified, within the iSPN/D2 cluster, a distinct yet small subcluster, here provisionally named ‘outlier D2’ (O-D2), which appeared in all samples across both phenotypes in both murine and human species. The O-D2 cluster is transcriptomically closer to the striosomal identity than to that of the matrix and shares markers with the ’D1/D2 hybrid’ recently identified in the non-human primate^20^. On the other hand, O-D2 is likely distinct from ’eccentric SPNs’^19^ (Extended Data Fig. 3), which seems a class of striatal interneuron according to our preliminary data (Pineda et al., in preparation). These cell types, identified by ACTIONet^20–34^, co-clustered perfectly atop each other between controls and the HD samples (magenta and cyan dots, Fig. 1A-C), affirming the consistency of cell-type annotations across phenotypes. There were no differences in quality control metrics between them or in the fraction of cells discarded (Methods), indicating that cell identities were not sufficiently perturbed by mutant huntingtin to confound annotation. Correspondence of the cell-type annotations was further supported by the well-matched fraction of each SPN subtype in the entire SPN populations across samples (Fig. 1D).

Of note is the profound depletion of SPNs found in the human HD, especially, of S-D2 SPNs and S-D1 SPNs, followed by M-D2 SPNs (Fig. 1D). This pattern indicated a clear preferential vulnerability of striosomal SPNs in HD, alongside the well-known D2-predominent dysregulation in this disorder. As expected from previous work, in samples from the HD model mice, SPN loss was negligible despite the disturbed transcriptional profiles of SPNs.

This S-M compartmental sub-clustering is the first to be documented in human snRNA-seq samples; the result prompted us to examined S-M markers for potential conservation across species. We took the ratio of each gene expression level (i.e., fold change, FC) between S-D1 and M-D1, to identify potential striosome markers in the D1 population as genes with differential expressions surpassing abs(log_2_FC) > 0.1 and p values (< 0.001) adjusted for false discovery rates (FDRs). We identified potential striosome markers in the D2 population with the same criteria in the comparison between S-D2 and M-D2. The overlap of marker genes between human and two rodent lines (BL6 as zQ175 controls, and CBA as R6/2 controls) showed that universal striosome markers, i.e., expressed more highly in striosomes both in D1/D2 populations, were more conserved than those detected only in either D1 or D2 populations (Fig. 1I). Similarly, matrix, D1 and D2 markers were better conserved when they were shared in D1/D2 or S/M populations (Extended Data Fig. 4). Over-represented gene ontology (GO) terms in each of four universal markers were found largely to be mapped into 9 categories of GO terms, related to development, cell adhesion, metabolism, migration, synapse/signaling, blood circulation, localization/transport, and organization (see Supplementary Table 2 for GO IDs included). As shown in Figure 1J, in all 9 groups of GO terms, the identities of mapped genes were partially shared across human and mouse, i.e., identified as markers in humans and one of rodent strains (magenta and pink) or both strains (dark cyan). Extended Data Fig. 5 show across- between species correlation of differential expression of S/M markers dependent on compartments (i.e., log_2_FC). The majority of striosome markers in one species were also expressed more highly in striosomes than in matrix in the other species (1st and 3rd quadrants of plots shown in Extended Data Fig. 5), but a noticeable number of markers reversed their compartmental preference from rodents to humans (2nd and 4th quadrants). For example, *CNTN5* is a striosome marker in human species, but a matrix marker in rodents. Overall conservation (i.e., overlaps) of gene identities were similar for D1/D2 markers (Extended Data Fig. 5), but D2 markers tended to be more conserved than D1 markers.

We found universal striosome, matrix, D1, and D2 markers to have distinctive patterns of GO enrichment (Fig. 1E-H). Across species, striosome markers (Fig. 1E) overrepresented development-related GO terms, whereas matrix markers (Fig. 1F) overrepresented those related to cell adhesion. In humans, D1 markers (Fig. 1G) overrepresented development- and migration-related GO terms, whereas D2 markers (Fig. 1H) overrepresented those related to synapse/signaling and blood circulation. These cell-type-specific patterns became further obvious when we included only universal marker genes conserved across both strains and mouse-human species for GO analysis (hyper-conserved markers, pink circles in Fig. 1E-H). The hyper-conserved markers include well-known key transcription factors involved in cell-type differentiation or signatures of every cell type after differentiation; *EphaA5*, *Htr2a*, *Oprm1*, and *Rxrg* were hyper-conserved striosome markers, *Epha4*, *Id4*, and *Zfhx3* were those for matrix, *Drd1*, *Ebf1*, *Foxp2*, *Pdyn*, *Reln*, and *Tac1* were those for dSPNs, and *Drd2*, *Oprd1*, and *Penk* were those for iSPNs. These patterns might indicate that, because the generation and maintenance of SPN cell-types are crucial for survival, mutant animals might have been eradicated through evolution if their mutations resided in the key marker genes irreplaceable for the differentiation or manifestation of SPN cell-types. The small number of surviving SPNs in human HD patients, even in the single Grade 1 case, rendered it impossible to analyze differentially expressed genes using the snRNA-seq data. We accordingly for our further analyses focused on the mouse models.

In each mouse model, we compared the expression of each transcript in HD mouse models with the level of expression in controls. We found 3,609 genes in R6/2 and 2,446 genes in zQ175 to be significantly upregulated or downregulated in the HD models in at least one cell type, as judged by the criteria of abs(log_2_FC) > 0.1, with p < 0.001. Figure 2A illustrates HD-associated alteration of Jensen-Shannon distances between each pair of cell types in transcriptional space for the two HD models (see Methods). Distances measured in the model mice were subtracted from those in controls. All resulting metrics (Fig. 2A) had negative values, indicating that the difference between transcriptomes for every pair of SPN types (e.g., M-D1 vs. S-D1) were lessened, i.e., they became more alike, in both zQ175 mice and R6/2 mice.

**Fig. 2.**
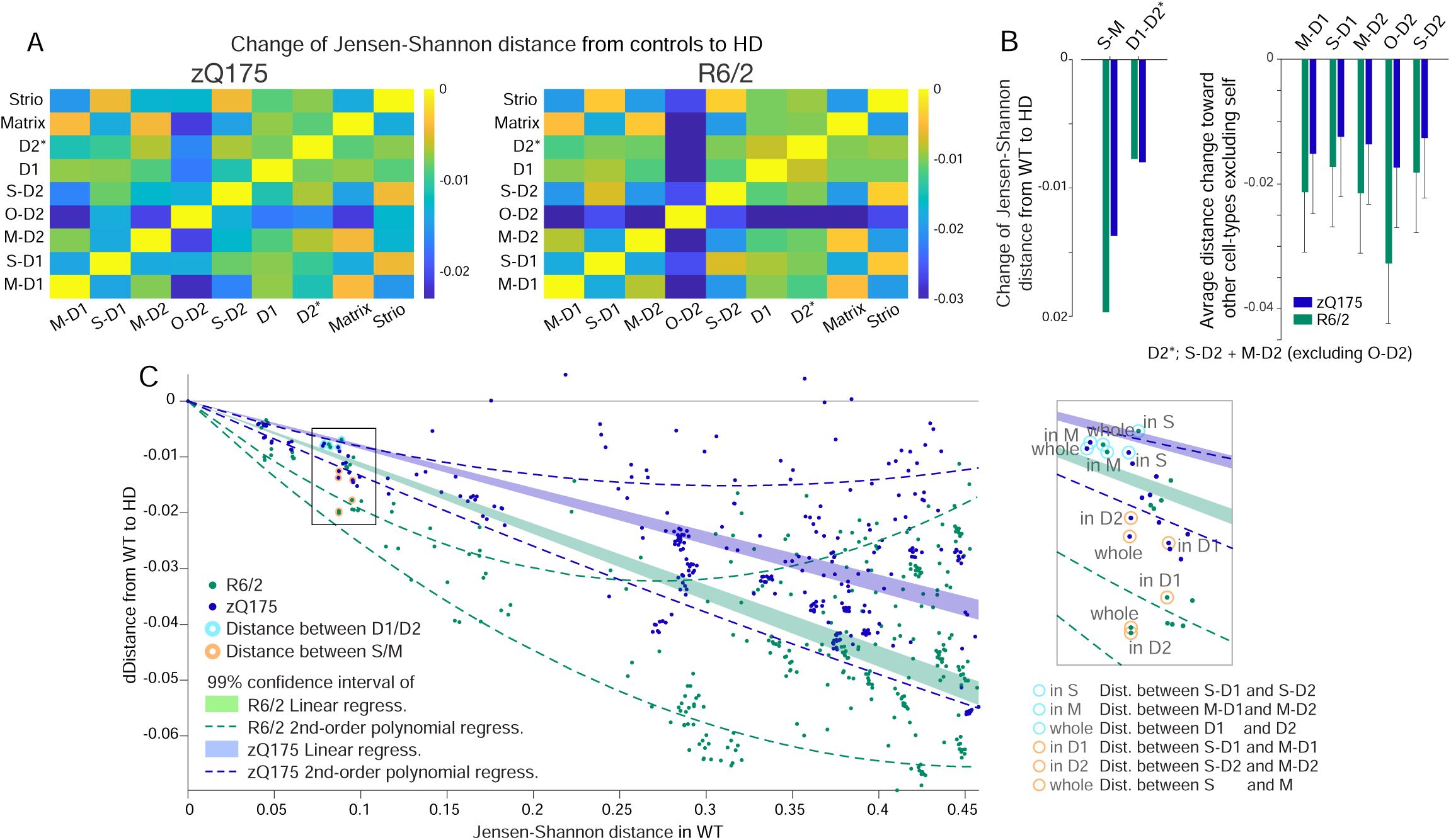
Striosome-matrix transcriptomic distinctions is more vulnerable than those of D1-D2 in HD. **A,** Jensen-Shannon distances between pairs of cell types for zQ175 (left) and R6/2 (right) as compared to control. Negative values indicate loss of transcriptomic distinction. **B**, Summary of loss of transcriptomic distinctions measured by Jensen-Shannon distance. Distinction between striosomes and matrix (S-M) and between dSPN and iSPN (D1-D2*) are shown. Right: Loss of each cell-type identity. Blue (zQ175) and green (R6/2) bars indicate averages. D2* does not include O-D2. Error bars indicate 95% confidence intervals. **C**, Loss of Jensen-Shannon distances in HD as a function of those in controls are shown for every pair of cell types in the striatum in zQ175 (blue) and R6/2 (green) models. Data points corresponding to S-M distinctions are demarcated by orange circle, whereas those for D1/D2 are demarcated by cyan circles. Loss of distance (i.e., loss of transcriptomic distinction, in HD) was larger for endogenously more distinct cell-type pairs in controls as captured by the 99% confidence intervals of linear regressions (shades) or second order polynomial regression (broken lines). Inset: enlarged image of boxed area. Note that loss of distance between S/M populations is larger than expected from the entire striatal dataset, whereas loss of distance between D1/D2 populations is smaller.

To our surprise, we found that this loss of distinctiveness was greater for the S-M axis of differentiation than for the direct-indirect pathway (D1-D2) SPN axis of differentiation in both HD models (the D2 population, setting aside the idiosyncratic O-D2 for future study (Pineda et al., in preparation; Fig. 2B). To judge whether the loss of distance between compartments was larger than expected from the distance patterns for the entire population of striatal cell-types, not only SPNs, as a control for the possibility that the loss of SPN distance measurements simply reflected such endogenous differences, we plotted the loss of Jensen-Shannon distance in the HD models as a function of endogenous distance between all possible pairs of cell types identified in the control striatum (Fig. 2C). Losses of distance between striosome and matrix (orange circles) were larger than expected from the linear or second-order polynomial regression of the entire striatal dataset; by contrast, losses of distance between D1 and D2 (cyan circles) were smaller than expected. These results indicate that, of the two classic axes of SPN classification in the striatum, the S-M axis is more imbalanced than the D1-D2 axis in the HD models, and demonstrate that this skewed abnormality coexists alongside the well-known preferential vulnerability of iSPNs in HD.

To probe for mechanisms that might lead to blurring of the transcriptomic distinctions along S-M vs. D1-D2 axes, we again focused on the marker genes and examined their upregulation or downregulation (Fig. 3). We identified universal or selective striosome markers as described above, respectively, as those transcripts for which expression was (1) significantly higher in striosomes than in matrix both in dSPN and iSPN populations in controls, or (2) significantly higher in either of these populations considered singly (S-D1 > M-D1, and/or S-D2 > M-D2, abs(log_2_FC) > 0.1 and p < 0.001). Similarly, we identified universal or selective matrix markers, respectively, as those transcripts for which expression was significantly higher in matrix than in striosomes in both or either in dSPN and iSPN populations in controls (S-D1 < M- D1, and/or S-D2 < M-D2). A consistent pattern emerged: striosome markers were more *up*regulated in matrix SPNs, but were more *down*regulated in striosomal SPNs, for both models. Conversely, matrix markers were more *up*regulated in striosomal SPNs, and more *down*regulated in matrix SPNs in both models. The upregulations and downregulations followed a gradient pattern from M-D1, M-D2, S-D1, S-D2, to O-D2 (Fig. 3A-I).

**Fig. 3.**
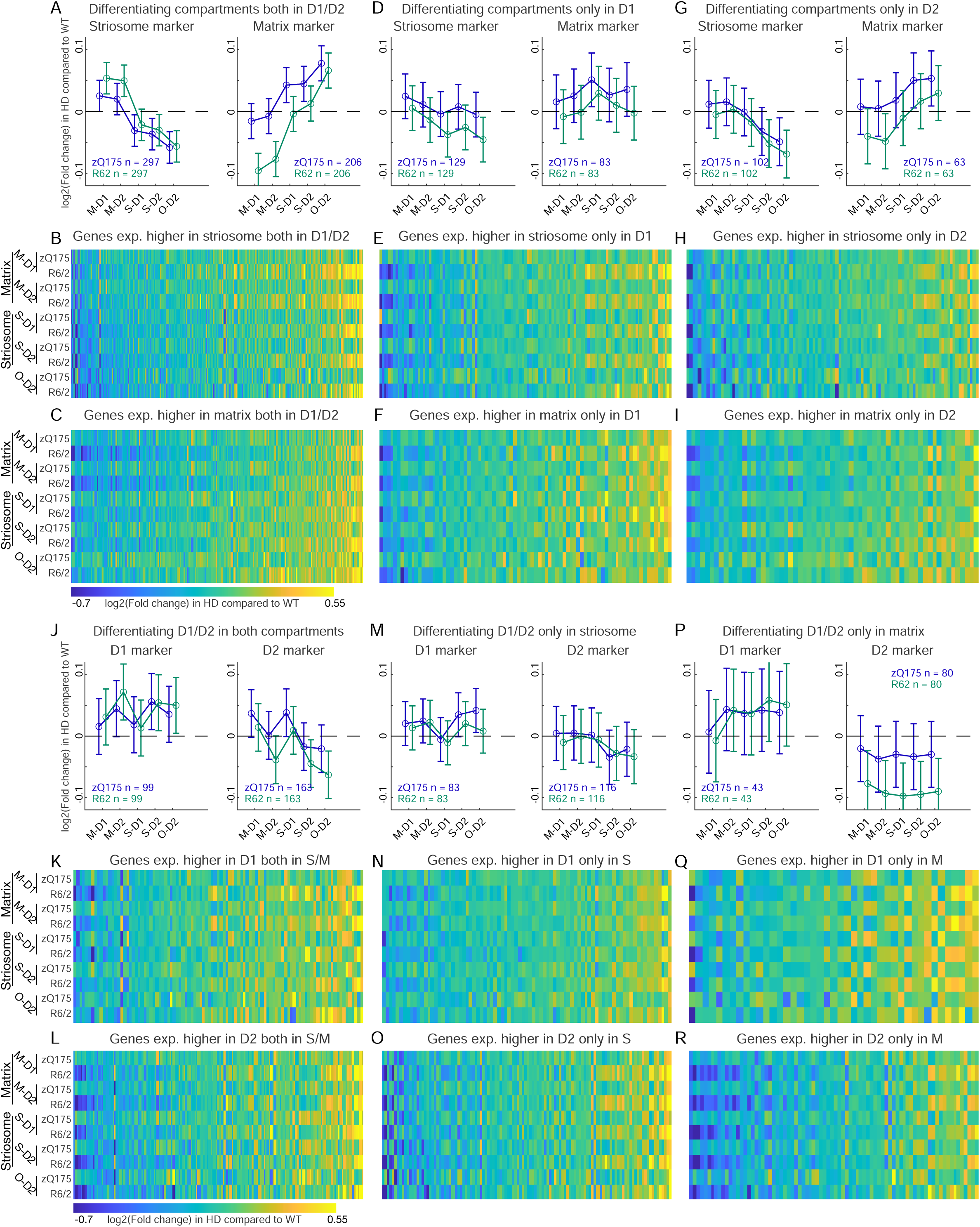
S-M markers but not D1-D2 markers exhibit cell-type-dependent dysregulation to blur their transcriptomic, discriminative identities. **A,** Alteration of striosomal (left) and matrix (right) marker expressions, which differentiated the control compartments both in dSPN and iSPN populations in zQ175 (blue) and R6/2 (green) models. Error bars indicate 95% confidence intervals. **B and C**, Alteration of striosome (**B**) or matrix (**C**) marker expressions in **A**. **D-I**, Same as in **A-C**, but for markers differentiating the compartments only in D1 (**D-F**) or D2 (**G-I**) population. **J**, Alteration of dSPN (left) and iSPN (right) marker expression, which differentiated D1-D2 in both compartments in controls, for each model. **K and L**, Alterations of D1 (**K**) or D2 (**L**) marker expressions in **J**, shown for each cell type of each model. **M-R**, Same as in **J-L**, but for markers differentiating D1/D2 only in striosomes (**M-O**) or in matrix (**P-R**). See also Extended Data Fig. 6.

These findings indicated that both SPNs in striosomes and SPNs in matrix exhibited a loss in their endogenous identities. This pattern of dysregulation, tending to cancel out differential expressions of S-M markers, was clear in those transcripts differentiating S-D2 and M-D2 (Fig. 3A,G) but was not uniformly detectable in those differentiating only S-D1 and M-D1 (Fig. 3D). The patterns held even after applying stricter criteria to define markers; we found similar, even clearer, patterns when we included S-M markers only if they differentiated compartments to the larger degree (abs(log_2_FC) > 0.2 rather than 0.1 and p < 0.001, Extended Data Fig. 6). This result indicates that the loss of S-M transcriptomic distinction is not due to the dissipation of weakly differentiating markers, but reflected the core pattern that the prime compartmental markers followed. The abnormality in iSPNs in these HD models, thus, included an obscuring of compartmental differentiation between S-D2 and M-D2.

In sharp contrast to the compartmental markers, D1-D2 markers did not alter their expression patterns so as to diminish their endogenous identities as dSPNs or iSPNs. Taking genes differentiating D1 and D2 in both or either compartment, D1 markers (S-D1 > S-D2, and/or M-D1 > M-D2, abs(log_2_FC) > 0.1 and FDR-adjusted p < 0.001) were inconsistently upregulated or downregulated across S/M and D1/D2 cell types. Thus, as groups of genes, significant upregulation in some genes cancel out the significant downregulation in others, rarely reached significance in either model (left panels in Fig. 3J,M,P). The D2 markers in the matrix of R6/2 mice, but not of zQ175 mice, were significantly downregulated irrespective of D1 or D2 cell types (right panel in Fig. 3P). Thus, in contrast to the clear and robust cell-type-specific dysregulation of S-M markers, dysregulation of D1-D2 marker expressions did not respect cell types, and thus maintained the distinction between D1-D2 populations, even though they distorted the profiles of genes distinguishing these two SPN classes.

Next, to identify cell-type-specific alterations, we examined the degree of dysregulation for each cell type. First, we included all marker and non-marker genes if they were dysregulated significantly in at least one cell type in the HD models (i.e., abs(log_2_FC) > 0.1 and p < 0.001). Absolute values of log_2_FC in each cell type (Fig. 4A) exhibited more severe patterns of dysregulation in R6/2 than in zQ175, confirming prior observations^17, 35^. It is of note, however, that our measurements were made in the context of different ages, i.e., comparing 9-week-old R6/2 mice to 6-month-old zQ175 mice.

**Fig. 4.**
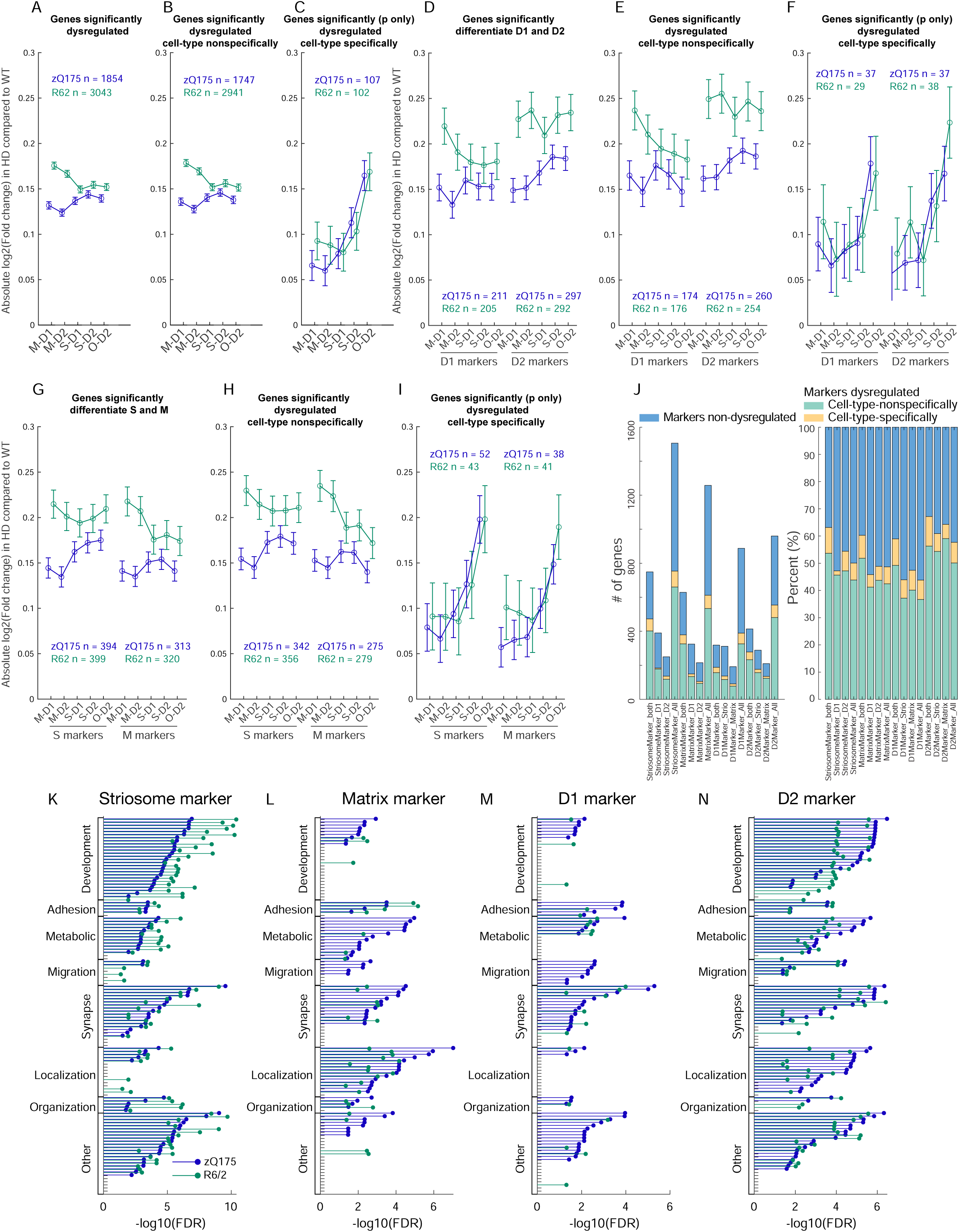
Cell-type-specific dysregulations reflect the intrinsic vulnerability shared across multiple HD models. **A-C,** Only cell-type-specific gene dysregulations are shared between the two HD models. Average degrees of dysregulation, i.e., absolute value of log2(fold change of the expression in HD as compared to that in controls), are shown for all dysregulated genes detected with the criteria of abs(log2FC) > 0.1 and p < 0.001 in at least one of four canonical cell types (**A**), or the subset of them which are unidirectionally dysregulated (i.e., upregulated, or downregulated in all four cell types, **B**). In **C**, we first selected genes with significant dysregula- tion (p < 0.001) in at least one of four canonical cell types, then further restrict to the genes that are dysregulated bidirectionally dependent on the cell types (i.e., upregulated in one cell type(s) and downregulated in another cell type(s)). Error bars indicate 95% confidence intervals. **D-F**, Same as **A-C** but restricted for D1-D2 markers. **G-I**, Same as **A-C** but restricted for striosome-matrix markers. **J**, Composition of patterns of dysregulation for D1-D2 markers and S-M markers. **K**, FDR for enriched GO terms in dysregulated striosome markers in zQ175 (blue) or R6/2 (green). Within the GO terms overrepresented in either of dysregulated striosome, dysregulated matrix, dysregulated D1, or dysregulated D2 markers, top (i.e., lowest FDR) 40 GO terms are included, and grouped into 9 categories. In each group, the GO terms are sorted by FDRs for zQ175. **L-N**, Same as **K** but showing FDRs of the enrichments in dysregulated matrix (**L**), dysregulated D1 (**M**), or dysregulated D2 (**N**) markers. See also Extended Data Fig. 7.

Second, we divided up the data depending on whether the dysregulation of a given gene was similar across all cell types or differed by cell type. For classification as being in the unidirectional, cell-type-nonspecific category (Fig. 4B), the transcript was upregulated or downregulated in all cell types with the requirement that the dysregulation was significant in at least one cell type (abs(log_2_FC) > 0.1 and p < 0.001). For categorization as bidirectional (Fig. 4C, i.e., cell-type-specifically dysregulated), the transcript was upregulated in some cell types and downregulated in other types, again with the requirement that the dysregulation be significant (p < 0.001) in at least one cell type. The cell-type-nonspecific dysregulations differed depending on which of the two HD models was examined, whereas cell-type-specific dysregulations were well aligned between the two, reflecting HD-related cell-type-dependent vulnerability held in common. The same rule was consistently observed when we included only D1-D2 marker genes (Fig. 4D-F) or only S-M marker genes (Fig. 4G-I), which were composed of similar proportions of genes dysregulated either cell-type-specifically or cell-type- nonspecifically (Fig. 4J). The shared pattern of dysregulation across the two models, measured as degree (i.e., absolute values of differences from their respective controls), had a hierarchy: it was highest in O-D2, followed by S-D2. This hierarchy thus reflected a multiplexing of the D2- dominant vulnerability with compartment-based vulnerability.

In order to obtain insight into biological pathways that might be especially dysregulated, we performed a GO analysis applied to (1) marker genes significantly dysregulated in at least one cell type (Fig. 4K-N), (2) cell-type-nonspecifically (i.e., upregulated or downregulated in all cell types) or specifically (i.e., upregulated in some cell types and downregulated in other types) dysregulated genes (Extended Data Fig. 7A-C), or (3) marker and non-marker genes dysregulated in each type of SPNs (Extended Data Fig. 7D-H). Consistent with the D2-dominant and striosome-dominant dysregulation, a wide range of GO terms were overrepresented in dysregulated D2 markers (Fig. 4N) and dysregulated striosome markers (Fig. 4K), especially those related to development. On the other hand, dysregulated D2 and matrix markers commonly overrepresented GO terms related to metabolic or localization processes. The differences in the patterns of enriched GO terms were more alike than those shown for universal S, M, D1, or D2 markers in Figure 1E-H, indicating that endogenous identities were defined by genes involved in distinct biological pathways, whereas in the HD models, dysregulations observed in the marker genes were involved in similar biological pathways. The data analysis thus uncovered nuanced, but considerable, differences in the biological pathways overrepresented in dysregulated S/M and D1/D2 marker genes.

To verify these transcriptional changes detected by snRNA-seq, we conducted fluorescence *in situ* hybridization (FISH) for the most prominent dSPNs and striosome markers in the two HD model mice (Fig. 5A-H). *Drd1* is downregulated in dSPNs of HD models (Fig. 5A,C,D), reflecting their loss of transcriptomic identities^36^. However, another strong marker of dSPNs, *Ebf1*, a well-known factor contributing to dSPNs differentiation^37, 38^, was upregulated in dSPNs (i.e., M-D1 and S-D1, Fig. 5A), but not in iSPNs, as if to rescue the loss of their identities, which was confirmed by FISH (Fig. 5C,D). *Nnat*, an imprinted gene implicated in the brain development^39, 40^, is a strong endogenous striosome marker and was downregulated in striosomes of the HD models (i.e., S-D1 and S-D2, Fig. 5E). In FISH data, its expression in the striosomes, but not in matrix, was significantly decreased, so as to lose the significant difference between the compartments in zQ175 (Fig. 5H). We found, in addition, that *Lypd1*, identified previously as a prototoxin that acts on nAChR as a snake neurotoxin^41^ and as one of markers of von Economo neurons^42^, was a robust striosome marker to identify striosomes in zQ175 mice, but it could not do so in R6/2 mice (Fig. 5G), indicating that markers even apparently robust in snRNA-seq data might alter their distribution patterns to hinder identification of compartmental identities by FISH.

**Fig. 5.**
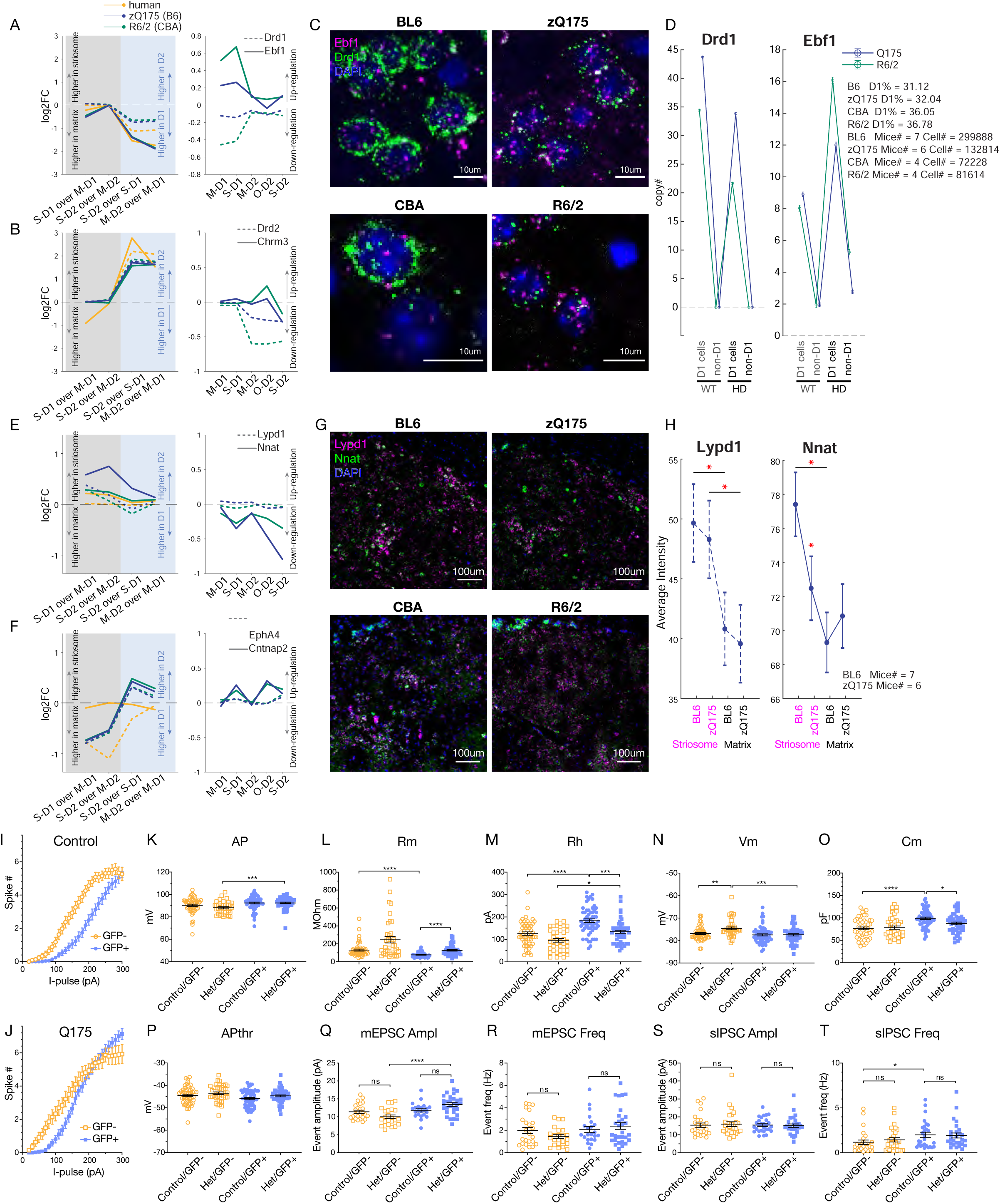
Histological and physiological loss of compartmental identity in HD model mice. **A,** snRNA-seq data for dSPN markers, i.e., Drd1 (broken lines) and Ebf1 (solid lines). Left: Differential expression was measured between the cell-type pairs indicated below and shown as log2(fold change). Right: Expressions in HD models are compared to those in controls. **B**, Same as in **A**, but for iSPN markers, i.e., Drd2 (broken lines) and Chrm3 (solid lines). **C**, FISH images for the two HD models compared to their controls, stained for Drd1 (green), Ebf1 (magenta), and DAPI (blue). **D**, Quantification of FISH image. Copy numbers (i.e., number of detected spots) for Drd1 (left) and Ebf1 (right) are shown separately for D1 and non-D1 cells in controls or HD models. Error bars indicate 95% confidence intervals. **E and F**, Same as in **A and B**, but for striosome markers (**E**), i.e., Lypd1 (broken lines) and Nnat (solid lines), or matrix markers (**F**), i.e., EphA4 (broken lines) and Cntnap2 (solid lines). **G**, Same as in **C** but for Lypd1 (magenda), Nnat (green), and DAPI (blue). **H**, Quantification of FISH image. Average intensity of FISH signals is shown for Lypd1 (left) and Nnat (right) separately for striosomes and matrix in zQ175 and in their controls. Error bars indicate 95% confidence intervals. * p<0.05. **I-T**, Using crosses of zQ175 and matrix-rich CalDAG-GEFI-GFP mice, electrophysiological properties of putative striosomal (GFP-negative, orange) and putative matrix (GFP-positive, purple) SPNs were examined ex vivo. Data are shown for zQ175 (squares) and control (circles) mice. Plots show current-frequency response for control (**I**) and zQ175 (**J**) mice, action potential amplitude (**K**) and threshold (**P**), membrane resistance (**L**), voltage (**N**) and capacitance (**O**), rheobase (**M**), mEPSC amplitude (***Q***) and frequency (**R**), and sIPSC amplitude (**S**) and frequency (**T**). *p < 0.05, **p < 0.01 and ***p < 0.001.

We also found *Chrm3*, coding muscarinic ACh receptor 3, to be as prominent an iSPN marker as Drd2 itself (Fig. 5B), and it was upregulated in O-D2 of the R6/2 (but downregulated in S-D2 of both models). The matrix marker, *Cntnap2*, an autism susceptibility gene^43, 44^, with as large compartmental difference in the expression as *Epha4*^45^ (Fig. 5F), was upregulated in SPNs other than matrix SPNs (S-D1, O-D2 and S-D2). Together, these results confirmed by histological evidence recapitulate the snRNA-seq pattern that disturbance of compartmental identities was greater than disturbance of D1/D2 identities in the HD models (Fig. 5A-H).

These transcriptional changes clearly raised the question of whether they could affect striatal function. As a first attempt to address this question, we conducted electrophysiological experiments in slice preparations aimed at assessing potential differences between striosomal and matrix cells (Fig. 5). We crossed zQ175 mice with CalDAG-GEF1-GFP mice differentially expressing GFP in matrix SPNs^46^. With whole-cell patch-clamp recordings, we replicated the higher excitability of putative striosomal (GFP-negative) SPNs as compared to putative matrix (GFP-positive) SPNs in control mice (Fig. 5I, n = 6). In zQ175 mice (n = 7), both types of SPNs increased their excitability as compared to their levels in controls (Fig. 5J), but the compartmental difference of excitability was lost in zQ175 mice. All significant dysregulations detected were cell-type-specific; membrane resistance (Fig. 5L), rheobase (Fig. 5M), and membrane capacitance (Fig. 5O) were significantly altered only in matrix (GFP-positive, shown in purple), whereas resting membrane potential was significantly elevated only in striosomal SPNs in the zQ175 (Fig. 5N). As a result, endogenous compartmental differences were either lost, as for measurements of membrane resistance (Fig. 5L), membrane capacitance (Fig. 5O), and sIPSC frequency (Fig. 5T), or were less significant, as for rheobase (Fig. 5M). Parameters related to the generation of action potentials and miniature excitatory and inhibitory potentials (mEPSCs and sIPSCs) also exhibited a few significant dysregulations in zQ175 SPNs.

## Discussion

Our findings demonstrate that the two canonical axes of striatal SPN organization, the direct (D1) and indirect (D2) pathway and the striosome-matrix compartmental subdivisions, are inter- dependently and differentially compromised in HD. The findings show that their identities categorized along the striosome-matrix (S-M) and dopamine receptor D1-D2 axes are multiplexed, yielding cross-axis vulnerabilities of these key projection neurons in both the R6/2 and the zQ175 HD mouse models. Our findings further demonstrate, surprisingly, that the transcriptomic distinctions along the striosome-matrix axis was more obscured than along the D1-D2 axis in both mouse lines. Echoing this pattern, in the Grade 1 HD human striatal snRNA-seq data set, the S-D2 sub-cluster was the most severely depleted of the entire SPN population, followed by S-D1 and M-D2. These consonant findings in the striatum of both mouse HD models and human HD suggest a profound differential susceptibility of both the S-M and D1-D2 canonical organizations known to govern the functional organization of the striatum. The consistent patterns across species further points to an excessive disturbance of the striosome-matrix organization and the excessive transcriptomic dysregulations of striosomes as phenotypes intermixed with the known vulnerability of indirect pathway D2-SPNs.

Our findings are constrained by limitations, some of which we mention here. The mouse samples were harvested from only a single timepoint; 9 weeks of age in R6/2 and 6 months of age in zQ175 models. We therefore cannot disentangle the time course of disease progression or the mechanistic sequence of pathologic events. Further, we were not able to examine subregions of the striatum from which the SPNs were sampled. This is a significant problem, given the well-known regional variations of molecular identities^7, 8, 18^ and the prototypical pattern of HD progression from the tail of the caudate nucleus forward in the human^47, 48^. We also could not fully analyze SPN transcriptomes in the human HD brain samples due to the limited numbers of surviving SPNs even in the Grade 1 HD case. This difficulty hampered our ability to forecast extension of the results for clinical translation. Nevertheless, the consistence of findings across human HD samples and two mouse HD models suggests that these limitations did not preclude the appearance of a common pattern of disease vulnerability. A potential link to the clinical findings is that in larger post-mortem samples examined by anatomical methods^13, 15^, differential vulnerability of striosomes has been found in HD cohorts from cases of early manifestation and cases of predominant mood disorder symptoms.

We further encountered in our analyses a group of D2-expressing putative iSPNs with an extreme and idiosyncratic transcriptional identity, and we here provisionally refer to them as outlier D2 (O-D2) cells. Their marker genes partially overlap with those of ’D1/D2 hybrid’ in primate^20^ and ’eccentric SPNs’ in mice^19^ (Extended Data Fig. 3). We treat the identity of O-D2 cells with caution in this study, in as much as they form a continuum not only with classical iSPNs but also with another non-SPN cell-type that expresses *Adarb2*, *Foxp2*, and *Olfm3* (Pineda et al., in preparation). The O-D2 category, although small, was notable in that we found, in the rodent HD models, transcriptomic dysregulations of this class of SPNs to be the most severe in the SPN population. However, in human HD samples, O-D2 was not the most affected population; the depletion of O-D2 cells in the Grade 1 HD samples was not as prominent as that of the S-D2 or S-D1 SPNs. Thus, we should be cautious about the mouse-human difference in the vulnerability of O-D2 cells, detected by different metrics, i.e., dysregulation magnitudes of transcripts in mice, and cell loss in humans.

The patterns of cell-type-dependent transcriptional changes that we identified were shared across SPNs in the two HD mouse models. Especially in genes dysregulated in opposite directions in different cell types, the cell-type-specific severities of transcriptional alterations were essentially identical in the R6/2 and zQ175 models. This finding supports previous observations^17, 49^ indicating that the transcriptomic dysregulations can be in common despite the distinct genetic makeup of these lines; the R6/2 model mice express an N-terminal exon1 fragment of mHTT, whereas the zQ175 mice express full-length mHTT. As a working hypothesis, we speculate that the cell-type-specific, model-invariant vulnerability stems from the cellular response to the mHTT exon1-like fragments, as a direct translation of exon1 transgene in R6/2 mice, or the products of the proteolysis of full-length mHTT protein translated in zQ175 mice. The hierarchical pattern of cell-type-dependent dysregulation, with highest levels in O-D2 followed by S-D2, was exhibited by all genes including non-markers, S-M and D1-D2 markers. The idiosyncratic transcriptomic responses exhibited by the two models were reflected in genes dysregulated in the same direction in all cell types.

It was between the cell-specific markers for the S-M and D1-D2 axes that the patterns of dysregulation were most clearly different in both mouse models. Within the S-M axis, the striosomal and matrix neurons each exhibited declines of their own markers but gains of the other’s markers, so that they became less differentiated from one another than in the control mice. Within the D1-D2 axis, by contrast, D1 and D2 markers were altered irrespective of cell type, without diminishing the transcriptomic D1-D2 distinctions, even though the distinctions were distorted. And the obscuring of the S-M axis was more severe in markers found in iSPNs than those found only in dSPNs (S-D1 vs. M-D1). These changes could contribute to the preferential abnormality of iSPNs in the HD model mice, and at the same time, emphasize the multiplexing of vulnerabilities across the D1-D2 and S-M axes. Potentially important regional distinctions across the striatum remain to be examined.

Our findings open a new view of the disturbance in balance between striosome-matrix and direct-indirect pathway circuits imposed by HD. Our findings open the possibility that these S-M and D1-D2 axes of striatal organization can be subject to distinct pathophysiological alterations. Among many possibilities for this intermixed, yet asymmetric vulnerability, we mention one of interest as it potentially links our findings to the clinical symptomatology of HD from pre-manifest to manifest stages. This is the working hypothesis that the S-M disturbance, possibly via a decrease of normal *HTT*, might precede at least in part the D1-D2 disturbance induced by the gain of *mHTT*.

Besides the gain of detrimental function of *mHTT*, loss of a beneficial function of wild-type huntingtin has been implicated in HD pathology^50–52^ and is gaining interest in response to early clinical failures of antisense candidates^53^. We do not here have direct data that allow us to distinguish effects of lowered levels of normal huntingtin from that of gain of *mHTT* in our samples, but our results are consonant with previous studies. Although full deletion of Htt is lethal, mice can live if the expression of a single allele of Htt is rescued at P21; they exhibit deficient compartmental organization and develop heterotopias expressing both matrix and striosome markers^54^. This abnormality in striosome-matrix compartmentalization was reported in the context of a homozygous, normal allele deletion during development; we here report compartmental dysregulation, even in the context of heterozygosity, which should be milder but one more akin to the condition of HD individuals, who are heterozygous for *mHTT*. Thus, loss of function could also, concomitant with the gain of *mHTT* function^52, 55^, hinder the differentiation of striosomal and matrix SPNs. Such evidence is unavailable for human, but studies using induced pluripotent stem cells derived from HD patients^56^ have demonstrated delayed differentiation and an increased pool of striatal progenitors. Abnormality has further been detected by MRI in non-juvenile CAG expansion carriers as young as 6 years of age^57^.

By influencing development, the loss of a normal *HTT* allele in HD heterozygous individuals could hinder the anatomical compartmentalization of the striatum as well as the differentiation of SPNs to acquire transcriptional identities as striosomes (mostly earlier born) or matrix (mostly later born). By contrast, preferential disturbance of D2-expressing iSPNs, known to be positively correlated with motor manifestation and vast transcriptomic alteration in iSPNs, is detected later, just around the age of manifest onset, under the control of CAG repeat length^2, 4, 5^. Such intermixed but temporally staged disturbance of the S-M and D1-D2 axes would align with the mood disorders in the pre-manifest stage, and motor disorder in clinically manifest stage, of HD. If so, the decline of transcriptomic differences between striosomes and matrix might exist far earlier than the manifestation of symptoms, even before birth. If the dysregulation of D1-D2 markers reflect the failure of compensatory mechanisms, or the response to somatically expanded CAG repeats^58^, then the dysregulations observed here might not be detectable early in life. These are among critical issues raised by our findings that need resolution in advancing HD therapeutic strategies.

## Supporting information

Suuplemental Table 1

Suuplemental Table 2

Suuplemental Table 3

Suuplemental Table 4

Suuplemental Table 5

## Acknowledgement

We thank Dr. Tomoko Yoshida, Blaise Clarke and Ara Mahar for their expert help with the histology, Cody Carter for help with mouse breeding, Henry F. Hall for his help with the laboratory, and Yasuo Kubota for help with manuscript preparation. This work was funded by the CHDI Foundation (A-5552 to A.M.G), the Saks Kavanaugh Foundation (to A.M.G.), the National Institutes of Health (R01 MH060379 to A.M.G.; R01 NS100802 to M.H.), the Nancy Lurie Marks Family Foundation (to A.M.G.), the Simons Foundation (SFARI 306140 to A.M.G.), the JPB Foundation (to H.L. and M.H.), the Kristin R. Pressman and Jessica J. Pourian ‘13 Fund (to A.M.G.), and Mr. Robert Buxton (to A.M.G.).

## Author Contributions

A.M.G., J.R.C., M.H., S.S.P. and M.K. planned and initiated the research; S.S.P. and H.L. performed the single-cell experiments; J.R.C. curated striosome-enriched gene list; S.S.P. performed the initial analysis to identify S-M sub-clusters, and performed further analysis; A.M. performed major analysis of these data in tight consultation with A.M.G., and with the help and guidance of M.H., S.S.P. and M.K.; A.M. and A.M.G. co-wrote the manuscript, with input from all authors.

## Methods

### Animals

All mouse husbandry and experimental procedures were conducted with the approval of the Massachusetts Institute of Technology Animal Care and Use Committee. Mice were housed under pathogen-free conditions, with food and water provided ad libitum on a standard 12-hr light/12-hr dark cycle. No procedures were performed on the mice prior to the outlined experiments. For all studies, littermate mice were group-housed, and male littermates were used at ages described in the Method Details and figure legends. Only male mice were used given HD model differences in phenotype progression between male and female mice. Mice were assigned to experimental groups based on their genotype (all mice were used), and as individual biological replicates. B6CBA-Tg(HDexon1)62Gpb/1J mice (CAG repeat length 160 ± 5; Jackson Laboratories stock # 002810) were used as R6/2, and B6J.zQ175DN (Jackson Laboratories stock # 370832) knock-in congenic C57BL/6J mice were used as zQ175 model. Replicate number per mouse group and sample size was as reported previously^17^.

### Human samples

Post-mortem caudate and putamen tissue samples of Grade 1 HD and matched unaffected controls were obtained from the NIH NeuroBioBank or the University of Alabama at Birmingham.

### snRNA-seq and analysis

Nuclear isolation was performed as described in Lee et al.^17^ (n = 62,487 nuclei across twelve unaffected control and Grade 1 HD caudate and putamen samples; n = 112,295 nuclei across fifteen mice: eight isogenic control and seven R6/2 model mice, all at 9 weeks of age; n = 63,015 nuclei across eight mice: four isogenic control and four zQ175DN model mice, all at 6 months of age; samples described in Supplementary Table 1). Droplet-based snRNA sequencing libraries were prepared using the Chromium Single Cell 3’ Reagent Kit v3 (10x Genomics, Pleasanton CA) according to the manufacturer’s protocol and sequenced on an Illumina NextSeq 500 at the MIT BioMicro Center (zQ175DN mouse samples) or a NovaSeq 6000 at the Broad Institute Genomics Platform (R6/2 mouse samples and human samples). FASTQ files were aligned to the pre-mRNA annotated Mus musculus reference genome version GRCm38 or human reference genome GRCh38. Cell Ranger v6.0 (10x Genomics, Pleasanton CA) was used for genome alignment and feature-barcode matrix generation.

We used the ACTIONet and scran R packages to normalize, batch correct, and cluster single-nucleus gene counts. Batch-corrected data were used as input to the archetypal analysis for cell type identification (ACTION) algorithm^20–34^ to identify a set of landmark cells or ‘archetypes’, each representing a potential underlying cell state. Using ACTION-decompositions with varying numbers of archetypes, we employed the ACTION-based network (ACTIONet) framework^20–34^ to create a multi-resolution nearest neighbor graph. A modified version of the stochastic gradient descent-based layout method was used in the uniform manifold approximation and projection (UMAP) algorithm^59^, to visualize the ACTIONet graph. A curated set of known cell-type-specific markers (Supplementary Table 1) was used to annotate individual cells with their expected cell type and assign a confidence score to each annotation, and network connectivity was used to correct low-confidence annotations. Multiple iterations of this process were performed to identify and prune low quality cells. At each iteration, we removed cells with high mitochondrial RNA content (> 5% for mouse and > 20% for human), abnormally low or high RNA content (relative to the distribution of its specific cluster with an initial global cutoff of 500 unique genes), ambiguous overlapping profiles resembling dissimilar cell types (generally corresponding to doublet nuclei), and cells corresponding to graphical nodes with a low k-core or low centrality in the network (generally corresponding to high ambient RNA content or doublet nuclei).

Cell-type-specific pseudobulk differential gene expression (DGE) analysis was performed using ACTIONet and limma^60^ for sufficiently abundant cell types using age (human), sex (human), and disease (human and mouse) phenotype as design covariates and gene-wise single-cell-level variance as weights for the linear model. Genes were considered differentially expressed if they had an FDR-corrected p-value < 0.001 and an absolute log_2_-fold change > 0.1 for that cell type relative to the normal control group or the reference cellular sub-type. To ensure that DGE results were reproducible and robust to differences in cell type abundance, we sampled with replacement equal numbers of mice/individuals and cells per mouse/individual for each cell type and repeated the pseudobulk analysis. Lastly, we repeated the analyses using DESeq2 as the model-fitting algorithm in lieu of limma to ensure replicability across methods. In all cases, DGE results were consistent, and we used the pseudobulk limma results for all downstream analyses.

To determine transcriptomic distance, we computed the average gene expression vector for each subtype and calculated the pair-wise Jensen-Shannon divergence (JSD) using the philentropy R package between all subtypes. The JSD is a measure of similarity between two distributions in the interval [0, 1] with 0 denoting two identical distributions. The Jensen-Shannon distance was defined as the square root of the JSD. For Figure 2, we used the difference of this distance between phenotypes relative to the control to determine the extent of transcriptional identity loss in HD, with more negative values suggesting greater loss of identity.

### GO (gene ontology) analysis

All GO analysis used the PANTHER overrepresentation test (Released 20210224)^61^ and Gene Ontology database DOI:10.5281/zenodo.5228828 Released 2021-08-18, FISHER test (http://geneontology.org/). We searched for over-represented ’biological pathway’ GOs, corrected with false discovery rate. We used custom reference lists which only contain genes that were detected at all in the experiment in a given species in a given comparison analysis, to correct any possible bias originated from the experimental and/or analytic procedures.

As shown in Figures 1 and 4, GO terms found to be over-represented were categorized into 9 groups: development, cell adhesion, metabolism, migration, synapse/signaling, blood circulation, localization/transport, and organization. We categorized GO terms based on the keywords that they include, namely, development, genesis, differentiation, growth or generation for the development-category, synapse, synaptic, membrane potential, action potential, signaling or signal transduction for synapse/signaling-category, adhesion for cell-adhesion-category, metabolic, catabolic, biosynthetic for metabolism-category, taxis, locomotion, motility, migration for migration-category, blood, vasodilation, circulation, or circulatory for blood-circulation-category, localization or transport for localization/transport-category, and organization or assembly for organization-category. Exact GO terms assigned to each category is listed in the TermMembers sheet in Supplementary Table 2.

### Histological analysis

Mice were deeply anesthetized and then transcardially perfused with phosphate-buffered saline (PBS) followed by 4% paraformaldehyde in PBS. Brains were post-fixed in the same fixative for 24 hr and cryoprotected in 30% sucrose for 48 hr at 4°C, and sectioned at 30 µm thick for immunohistochemistry, and 10 µm for FISH using microtome. For immunohistochemistry, sections were blocked in 3% H_2_O_2_ prepared in PBS for 10 min, then blocked in TSA blocking reagent (Akoya biosciences, NEL744001KT), then anti-Mu opioid receptor antibody (abcam, ab134054, 1:5000) was applied to be incubated for 24 hr at 4°C. On day 2, secondary antibody (polymer HRP conjugated goat anti-rabbit antibody, Invitrogen, B40962) was applied then incubated for 45 min. Then sections were incubated in TSA plus Cy3 (Akoya biosciences, NEL744001KT) for 10 min. After that, Necab1 antibody (Sigma, HPA023629, 1:500) and somatostatin antibody (Millipore, MAB354, 1:100) were applied to be incubated for 72 hr at 4°C. On Day 3, secondary antibodies (goat anti-rabbit 488, Invitrogen A11034 and goat anti-rat 647, Invitrogen A21247, 1:300) were applied to be incubated for 2 hr at room temperature. Then sections were incubated in DAPI (Invitrogen, 62248, 1:1000), mounted on slide and cover slipped with ProLong Gold (Invitrogen, P36930). For FISH, we used RNAscope® Fluorescent Multiplex Detection Reagent (ACD Bio, 320851) to detect Nnat (432631), Lypd1 (318361-C3), Drd1 (461901-C2), Ebf1 (433411), and Chrm3 (437701-C3) following the manufacturer’s protocol.

### Slice preparation

Mice were euthanized by decapitation under isoflurane anesthesia. The brain was rapidly removed and cooled in ice-cold oxygenated cutting N-methyl-D-glucamine (NMDG) solution composed of (in mM): NMDG 105, HEPES 20 mM, KCl 2.5, glucose 5, CaCl_2_ 0.5 MgSO_4_ 10, NaH_2_PO_4_ 1.2, NaHCO_3_ 26, sodium pyruvate 3, sodium ascorbate 5, thiourea, buffered to pH 7.4 with HCl. Parahorizontal slices (300 µm) were prepared using a vibratome (Leica VT1000S; Leica Microsystems) and allowed to recover at 32°C inoxygenated cutting NMDG solution for 10 min. After this recovery period, slices were transferred to a holding chamber containing normal ACSF composed of (in mM): NaCl 124, KCl 3.5, NaH_2_PO_4_ 1.2, NaHCO_3_ 26, Glucose 11, MgSO_4_ 1.3, CaCl_2_ 2.5, (pH 7.3-7.4, osmolarity ∼300 mOsm), and allowed to recover for 1 hr at room temperature prior to electrophysiological recordings.

#### Membrane properties

Whole-cell patch-clamp recordings from visually identified SPNs were performed in the dorsal striatum at room temperature using fire-polished pipettes (5-7 mW) containing (in mM): K-gluconate 105, KCl 30, EGTA 0.3, HEPES 10, MgCl_2_ 4, Na_2_ATP 4, Na_3_GTP 0.3, Tris-phosphocreatine 10, pH adjusted to 7.2 with KOH. Data were collected from both GFP-positive and GFP-negative cells for each genotype using standard GFP fluorescence excitation and emission filters. Selected membrane properties (Vm, Rm, Rh, Cm, AP amplitude, AP threshold) were recorded. Rm and Cm values were collected using the automated “membrane test” function (+5 mV step; Vh = −80 mV). Rheobase (Rh) was determined using 300 msec current pulses (10-pA steps) from resting membrane potential, sampled at 10 KHz. Firing frequency curves were derived from spike number evoked during 10 pA depolarizing steps (300 msec).

#### mEPSCs and sIPSCs

For mEPSCs, whole-cell patch-clamp recordings were made with pipettes containing (in mM): Cs-methansulfonate 110, EGTA 10, HEPES 10, TEA-Cl 10, NaCl 14, CaCl_2_ 1, Mg-ATP 5, Na_2_GTP 0.5, Qx314-Cl 5, pH 7.2. mEPSCs were isolated by including 0.5 µM TTX and 40 µM picrotoxin in the bath solution, filtered at 1 KHz, and collected continuously for 5 min at a holding potential of −80 mV. For spontaneous IPSCs (sIPSCs), whole-cell patch-clamp recordings were made with pipettes containing the same solution described for mEPSCs (Erev_Cl_ = −38 mV). sIPSCs were isolated by including 5 µM CNQX in the bath solution, filtered at 1 KHz and collected continuously for 5 min at a holding potential of −80 mV.

### Transcriptional profiling

Use of the relevant pipelines for quantification, determination of normality of data, and appropriate statistical analysis of snRNA-seq data are as described previously^17^. Data met assumptions of the statistical approach based upon the experimental design in each case. Differential gene expression analysis of the snRNA-seq data was performed on a by-cell-type basis using both limma and DESeq2 in order to independently confirm statistical results.

As defined in the main text, expression differences were considered significant if they had an abs(log_2_FC) > 0.1, with FDR-adjusted p < 0.001.

### Data analysis of FISH data

We used HALO (indica labs, v3.3.2541.262) to analyze data taken by TissueFAXS Whole Slide Scanning System from TissueGnostics (Zeiss 20x 0.5 NA EC Plan-NEOFLUAR objective, Hamamatsu Orca Flash 4.0 V2 cooled digital CMOS camera C11440- 22CU for fluorescence imaging, Lumencor Spectra X light engine, motorized stage). To prepare for semi-automated image analysis, we exported all images taken by TissueFAXS systems in 16- bit format with the full range of bit color intensity values (0-65536).

For the analysis shown in Figure 5C,D, we applied customized algorithm modified from Indica_Labs_-FISHIF v2.1.5 to data of dorsal striatal regions (anterior, mid, and posterior sections from each animal). Briefly, we set threshold (including contrast threshold, intensity, segmentation aggressiveness, size and roundness) to detect nuclei by using DAPI channel, and marked surrounding cytoplasm. For Drd1 and Ebf1, we individually optimized contrast threshold, intensity, spot size, and segmentation aggressiveness to detect and count each copy of mRNA signals. We then extracted copy numbers of Drd1 and Ebf1 in individual cells identified from DAPI signals and define dSPNs as cells with Drd1 copy # > 4. We tried different threshold of Drd1 copy # to define dSPNs (e.g., 2, 6, 10), to confirm the results are largely the same.

For the analysis shown in Figure 5G,H, we applied customized algorithm modified from Indica_Labs_-Area Quantification FL v2.1. Briefly, we manually scored Lypd1 channel of each section (dorsal striatal regions at anterior, mid, and posterior levels from each animal), to annotate striosome areas (i.e., regions of interest, or ROIs) as Lypd1-positove areas. As matrix ROIs, we copied the size and shape of the nearby corresponding striosome ROI, and pasting the ROI to the surrounding Lypd1-negative region. We then measured the average intensities of Nnat and Lypd1 in the striosome and matrix ROIs.

### Data analysis of *ex vivo* experiments

All data were analyzed with Clampfit (Molecular Devices) and/or customized MATLAB routines. mEPSCs and sIPSCs were analyzed with Minianalysis software, using automatic event detection followed by visual verification. Both genotypic and GFP-positive/negative comparisons were made using a non-parametric ANOVA (Kruskal-Wallis) followed by a Dunn’s post hoc test. Firing frequency curves were analyzed by 2-way ANOVA. Data are presented with mean ± SEM.

### Data and code availability

All sequencing datasets generated as part of this study are publicly available in NCBI GEO under accession # GEO: GSE152058. Code used to construct ACTIONet and identify cell types is accessible from https://github.com/shmohammadi86/ACTIONet.

## Supplementary Tables

**Supplementary Table 1. List of samples used in this study, the curated set of compartmental markers, and list of O-D2 markers, related to Figure 1.** We searched for genes whose expression in O-D2 is highly enriched as compared to the ground average expression level in the striatum (including non-SPNS), i.e., abs(log2FC) > 0.1; p < 1e-05, and not enriched in any other cell types, i.e., abs(log2FC) < 0.1; p > 0.001.

**Supplementary Table 2.** List of GO terms enriched in universal striosome, matrix, D1, or D2 markers compared across human, BL6, and CBA samples, list of GO terms in the order shown in Figure 1E-H, and list of GO terms in the order shown in Figure 4K-N, related to Figures 1 and 4.

**Supplementary Table 3. Matrix of Jensen-Shannon distance between SPN subtypes, and matrix of Jensen-Shannon distance between all striatal subtypes, related to Figure 2.**

**Supplementary Table 4.** Dysregulation of all significantly dysregulated genes (i.e., including non-markers), significantly dysregulated striosome or matrix markers, and significantly dysregulated D1 or D2 markers are shown in separate sheet for each of R6/2 and zQ175 models, related to **Figure 4**. Within each sheet, genes dysregulated in a cell-type nonspecific manner (i.e., upregulated in all 4 canonical SPN subtypes, or downregulated in all 4 canonical SPN subtypes, with criteria abs(log2(fold change)) > 0.1 and FDR-adjusted p < 0.001), or specific manner (i.e., upregulated in one subtype while downregulated in other(s), with criteria FDR-adjusted p < 0.001) are listed in leftward columns as well.

**Supplementary Table 5.** List of GO terms enriched in genes dysregulated significantly at least in one cell-type (Any), cell-type-nonspecifically dysregulated genes (NONspe), and cell-type-specifically dysregulated genes (Spe), separately for zQ175 and R6/2 model, related to Extended Data Fig. 7. **For the categorization, see main text.**

---

**Extended Data Fig. 1.**
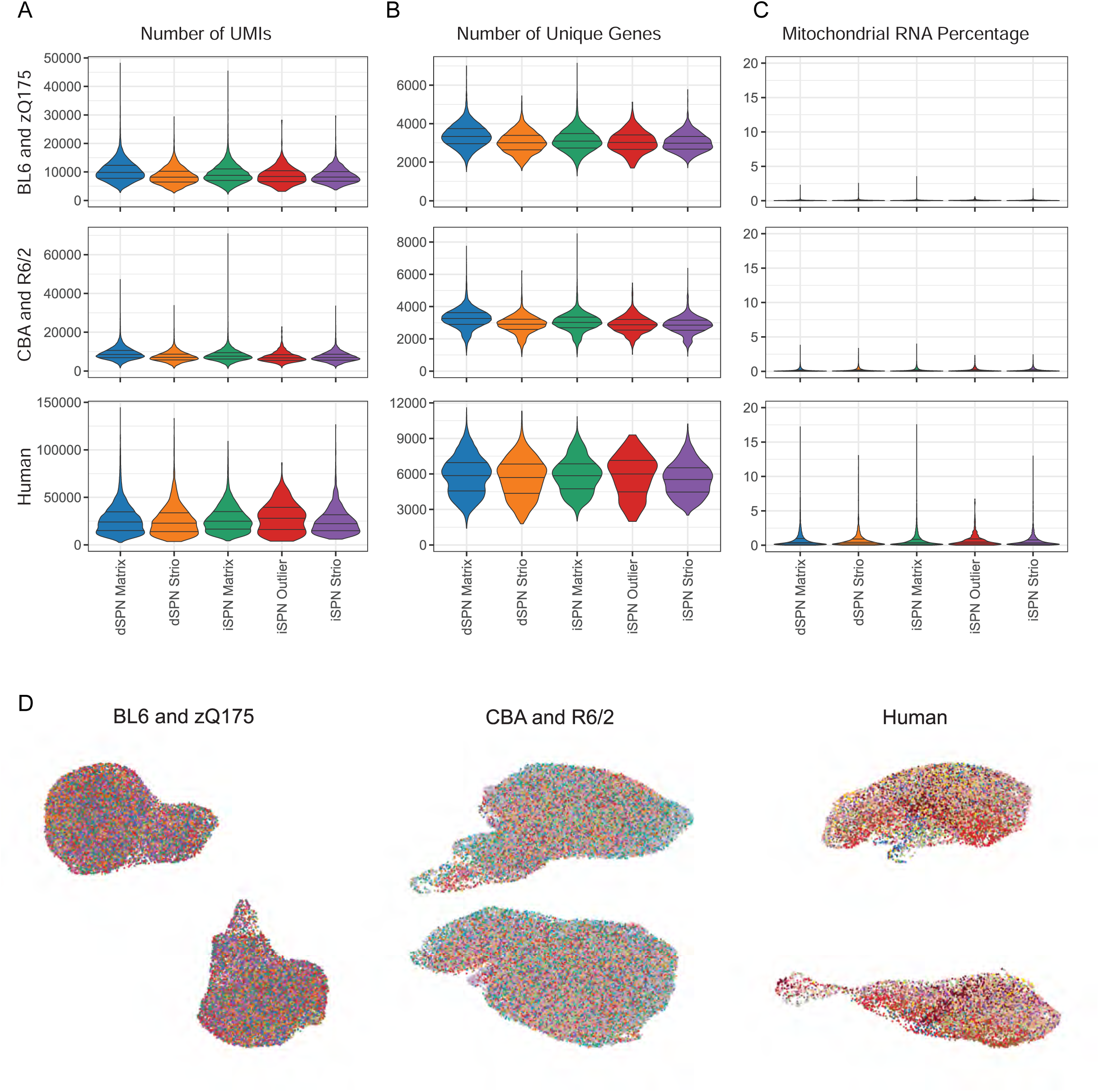
Benchmarking of snRNA-seq and data analysis by ACTIONet in this study. **A-C**, Quality control metrics of snRNA-seq data, with the number of unique molecular identifiers (UMIs; i.e. unique transcripts, A), number of unique genes (**B**), and percentage of mitochondrial RNA (**C**) per cell type. **D**, ACTIONet UMAP plots shown in Fig. 1 colored by individual samples.

**Extended Data Fig. 2.**
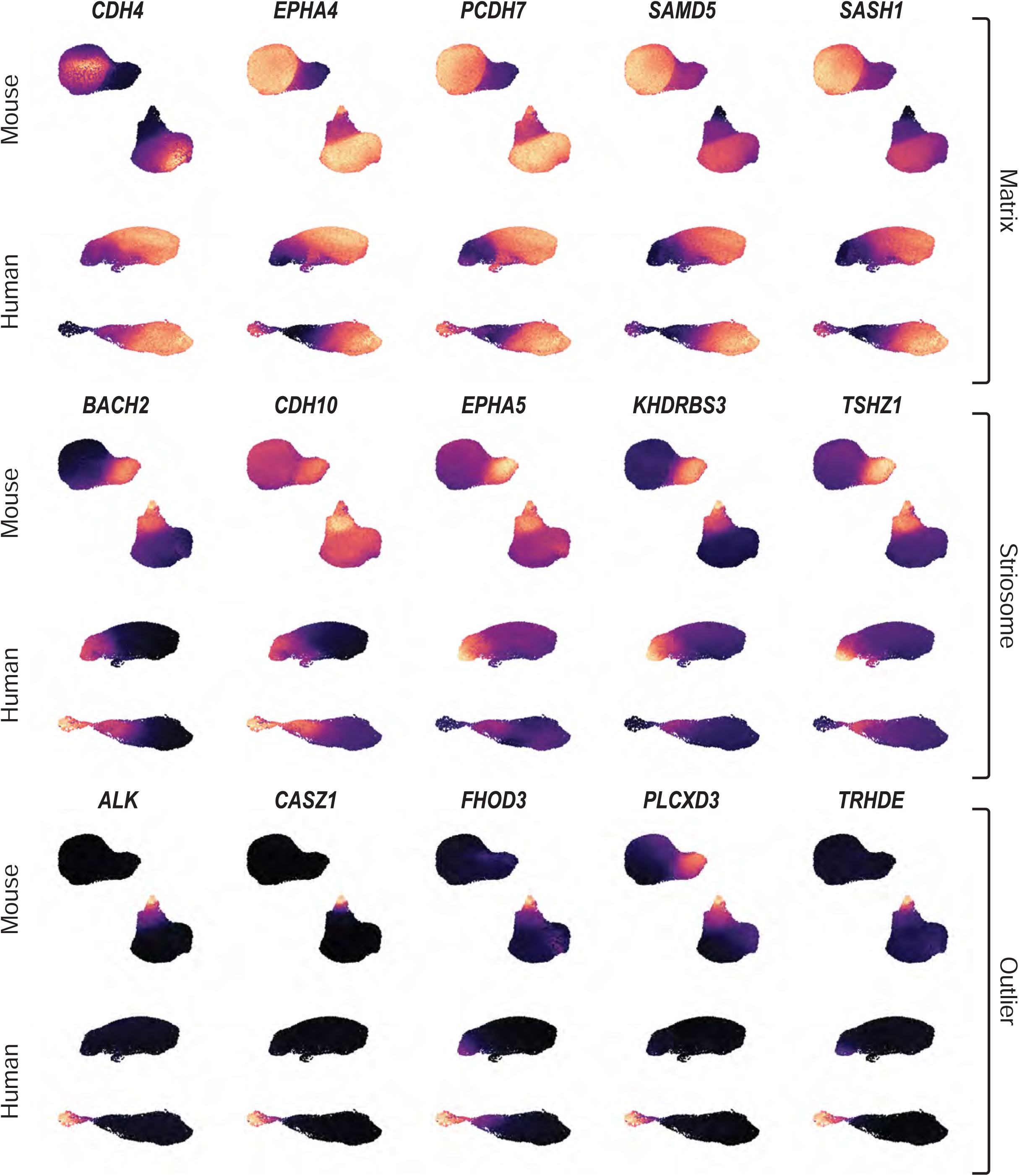
Expression of cell-type specific markers. Relative expression of representative matrix, striosome, and iSPN outlier marker genes in human and mouse (BL6 and zQ175) snRNA-seq datasets shown in Fig. 1.

**Extended Data Fig. 3.**
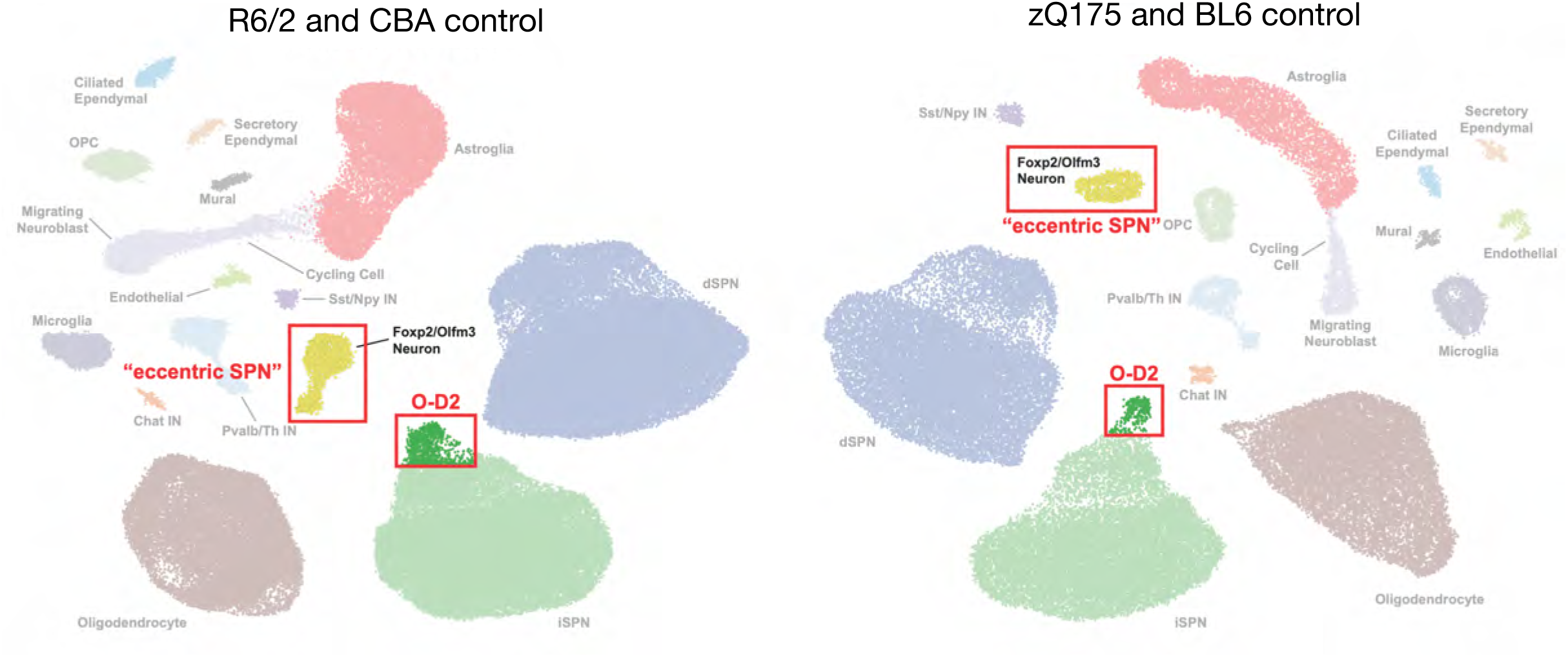
Relative relationship of O-D2 and eccentric SPNs on the ACTIONet plot. Two-dimensional ACTIONet graphs including all cell types in the striatal samples from R6/2 (left) and zQ175 (right) mice. Note that Olfm3-cells, i.e., previously named “eccentric SPNs” (Saunders., et al., 2018), form a distinct cluster separate from SPNs, whereas Outlier-D2 cells form a subcluster within the parent D2 SPN cluster (from Figure 4 in Lee et al., 2000).

**Extended Data Fig. 4.**
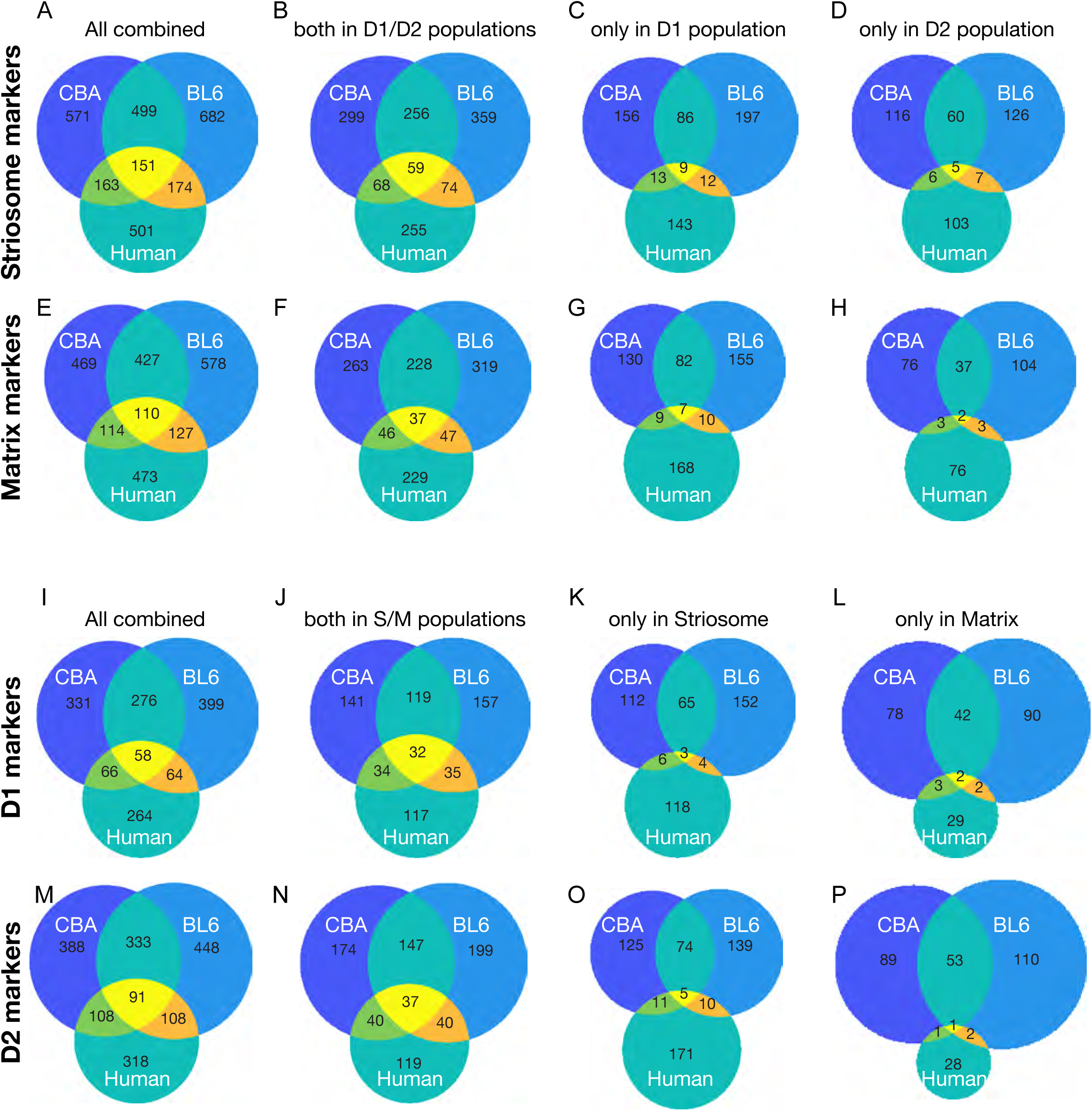
Genes differentiating S/M in both D1/D2 populations are more conserved than those differentiating S/M only in either population. (**A-D**) Venn charts showing the shared and unique striosome markers (i.e., expressed higher in striosomes than in matrix), detected in D1 and D2 population (**B**), only in D1 population (**C**), only in D2 population (**D**), or listed as either one of the three (**A**). (**D-H**) Same as **A-D**, for matrix markers (i.e., genes expressed higher in matrix than in striosomes). (**I-L**) Same as **A-D**, for D1 marker (i.e., genes expressed higher in D1 than in D2 cells). (**M-P**) Same as **A-D**, for D2 marker (i.e., genes expressed higher in D2 than in D1 cells). Note that universal markers, i.e., S/M markers that are consistently detected in D1 and D2 population (**B, F**) or D1/D2 markers that are consistently detected in both compartments (**J, N**), are more conserved than markers differentiating only in a specific SPN population.

**Extended Data Fig. 5.**
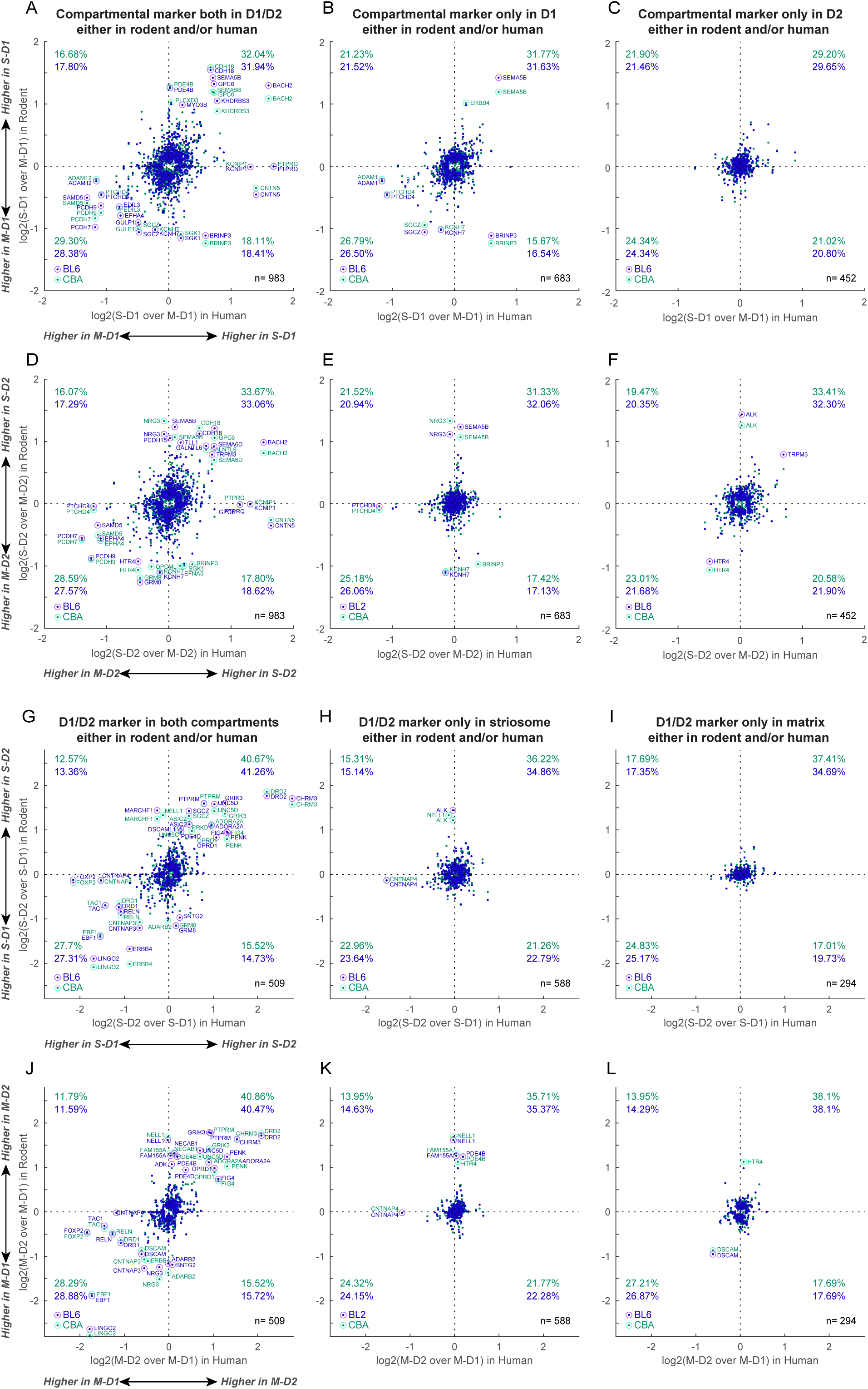
Although majority of S/M and D1/D2 marker genes are conserved across species, notable number of markers inverted their preference between rodents and human. (**A**) For universal S/M marker genes found either in rodents or human, we plot log2(fold change) of their expression in S-D1 as compared to that in M-D1. Data for BL6 (blue) and CBA (green) controls are shown. Percentage in each quadrant indicate the proportion of genes falling in that quadrant. (**B** and **C**) Same as **A**, but for compartmental markers found only in D1 (**B**) or D2 (**C**) population of either rodents or human. (**D**) Same as **A**, but showing log2(fold change) of expressions in S-D2 as compared to M-D2. (**E** and **F**) Same as **D**, but for compartmental markers found only in D1 (**E**) or D2 (**F**) population of either rodents or human. (**G**) Same as **A**, but for universal D1/D2 marker genes found either in rodents or human. (**H** and **I**) Same as G, but for D1/D2 markers found only in striosomes (**H**) or matrix (**I**) of either rodents or human. (**J**) Same as **G**, but showing log2(fold change) of expressions between M-D1 and M-D2. (**K** and **L**) Same as J, but for D1/D2 markers found only in striosomes (**K**) or matrix (**L**) of either rodents or human. We applied the criteria (abs(FC) > 0.1, p < 0.001) to define S/M markers that have a significant difference in the expression in striosomes and in matrix. Note that genes are included if they are listed as either of rodent (BL6 or CBA) or human markers. Thus, some genes appear in multiple panels; for example, KCNH7 is a universal striosome marker in rodents, whereas its differential expression across compartments was significant only in D1 population in human, appearing both in panels **A** and **B**.

**Extended Data Fig. 6.**
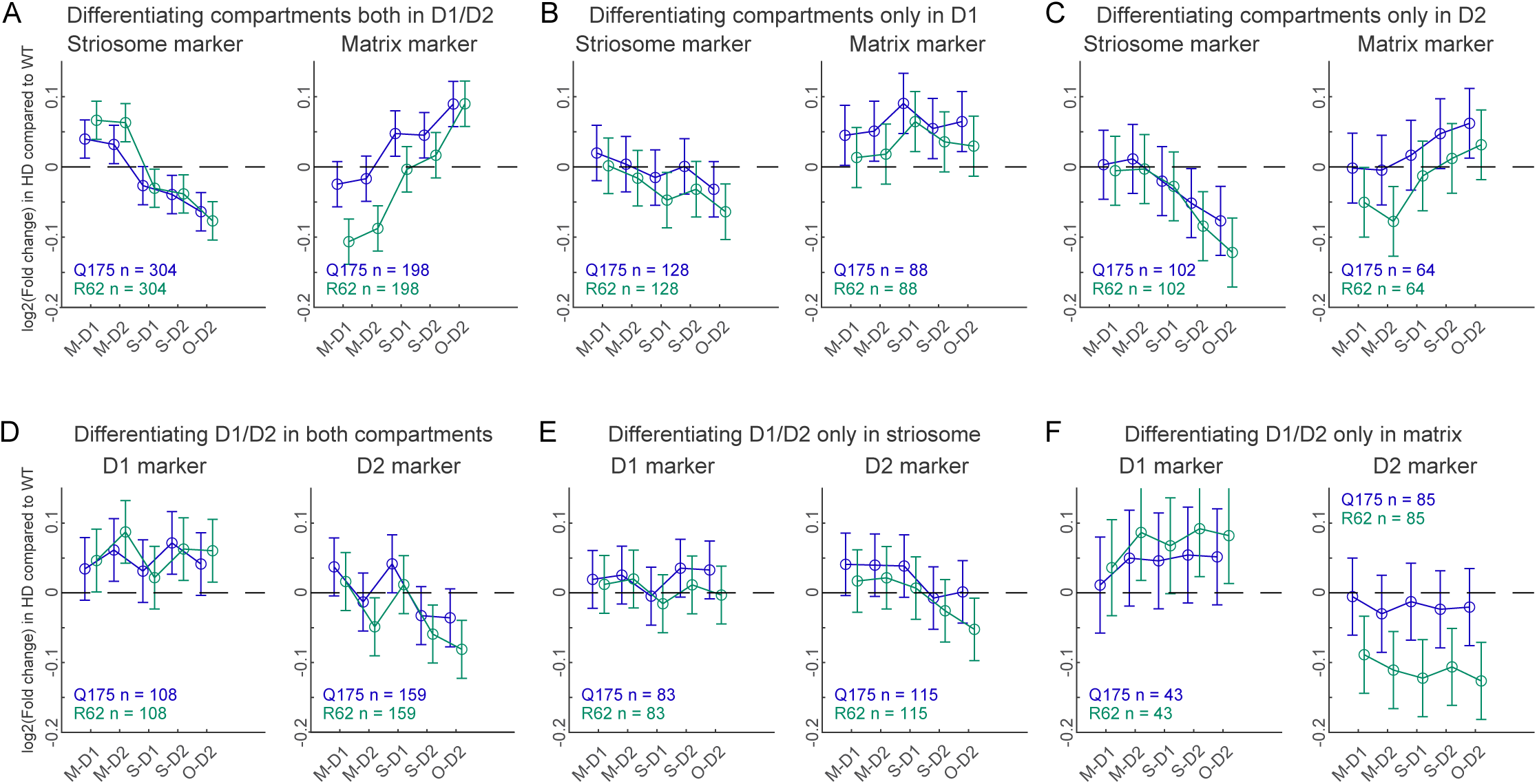
Same analysis as in Figure 3 but only including strong, highly differentiating markers. We applied more strict criteria to define as marker (abs(log2FC) > 0.2) than those used in the analysis described in the main text (abs(log2FC) > 0.1). The criteria of FDR adjusted p values were the same, i.e., p < 0.001. (**A**) Alteration of striosome (left) and matrix (right) marker expression, which is expressed more highly in striosomes or matrix both in D1/D2 populations, respectively, is shown for zQ175 (blue) and R6/2 (green) mice. Error bars indicate 95% confidence intervals. (**B** and **C**) Same as in **A**, but for markers differentiating S/M only in dSPNs (**B**) or in iSPNs (**C**). (**D**) Alteration of dSPN (left) and iSPN (right) marker expression, which is expressed more highly in D1 or D2 in both compartments, respectively, is shown for each model. (**E** and **F**) Same as in **D**, but for markers differentiating D1/D2 only in striosomes (**E**) or in matrix (**F**).

**Extended Data Fig. 7.**
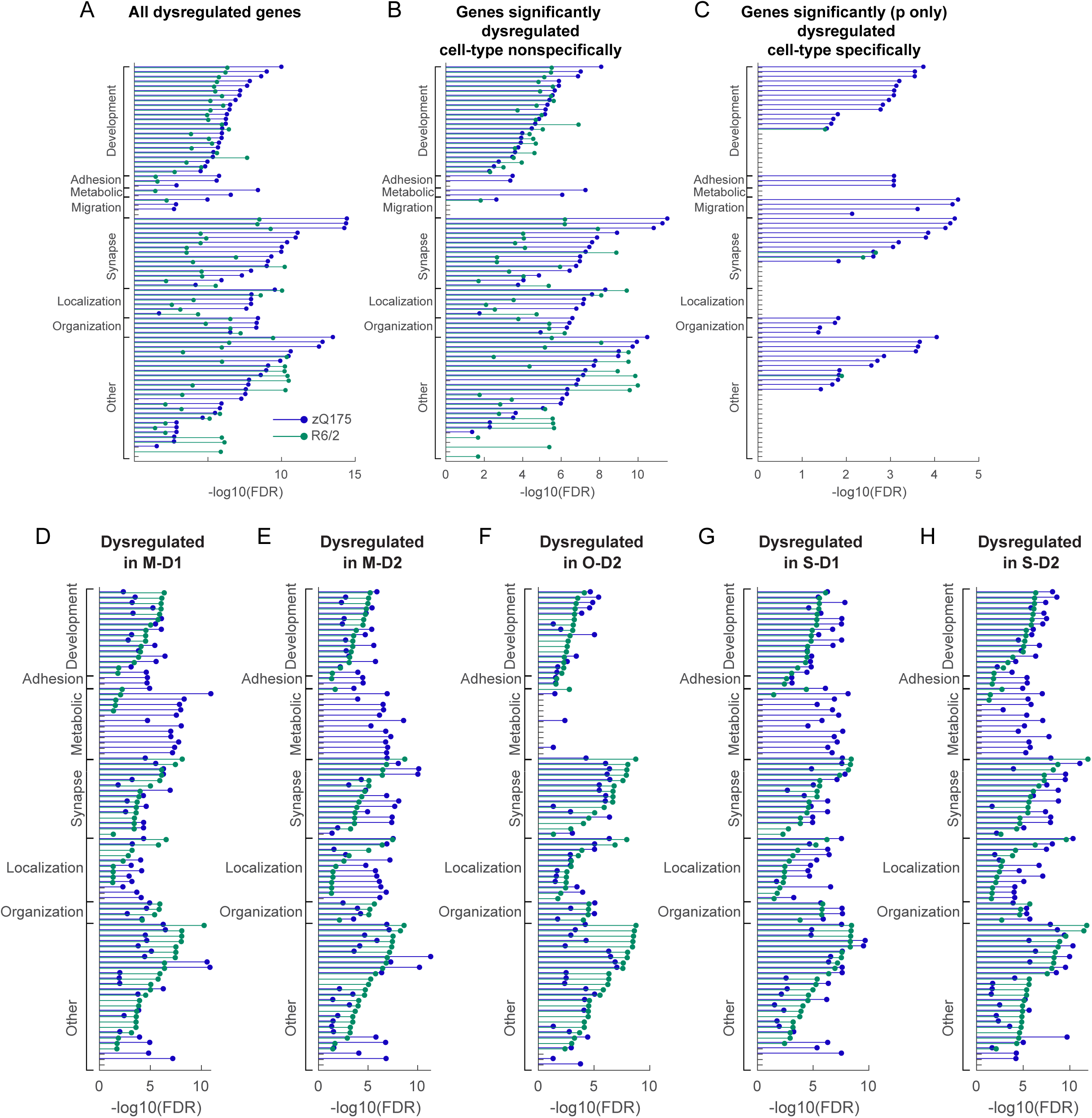
GO analysis on dysregulated genes. (**A-C**) FDRs for enriched GO terms are shown for all dysregulated genes detected with the criteria of abs(log2FC) > 0.1 and FDR-adjusted p value < 0.001 in at least one of four canonical cell types (**A**), or the subset of them that are unidirectionally dysregulated in all four canonical cell types (i.e., upregulation or downregulation in all four cell types, **B**). In **C**, we first selected genes with significant dysregulation (p < 0.001) in at least one of four canonical cell types, then further restricted to the genes that are dysregulated bidirectionally dependent on the cell types (i.e., upregulated in one cell type and downregulated in another cell type). (**D-H**) FDRs for enriched GO terms are shown for all dysregulated genes detected in M-D1 (**D**), M-D2 (**E**), O-D2 (**F**), S-D1 (**G**), or S-D2 (**H**). See Supplementary Table 5 for ID and full list of GO terms.

